# Compulsive drinking is associated with neural activity patterns reflecting diminished behavioral control and enhanced seeking representations in dorsal medial prefrontal cortex

**DOI:** 10.1101/2021.03.15.435169

**Authors:** Nicholas M. Timme, Baofeng Ma, David Linsenbardt, Ethan Cornwell, Taylor Galbari, Christopher Lapish

## Abstract

Drinking despite negative consequences (compulsive drinking) is a central contributor to high-risk alcohol intake and is associated with poor treatment outcomes in humans. We used a rodent model of compulsive drinking to examine the role played by dorsal medial prefrontal cortex (dmPFC), a brain region involved in maladaptive decision-making in addiction, in this clinically critical phenomenon. We developed novel advances in principal component and change point analyses to dissect neural population representations of specific decision-making variables. Compulsive subjects showed weakened representations of behavioral control signals that relate to drinking within a trial, but strengthened session-wide seeking state representations that were associated with drinking engagement at the start of each drinking opportunity. Finally, chemogenetic-based excitation of dmPFC prevented escalation of compulsive drinking. Collectively, these data indicate that compulsive drinking is associated with alterations in dmPFC neural activity that underlie diminished behavioral control and enhanced seeking.

## Introduction

Continued use of alcohol despite negative consequences (referred to as compulsive^1-3^) characterizes an advanced stage of addiction associated with poor treatment outcomes. Developing novel ways to treat addiction at this stage requires the identification of the brain mechanisms that underlie the decision to compulsively use alcohol. Isolating these mechanisms is difficult as decision-making deficits can exist both as a risk factor for and consequence of addiction^4^. Towards this goal, the current study assessed the decision to drink alcohol in a rodent model of compulsive drinking.

Impairments in decision-making are observed following excessive alcohol exposure^4^ and are accompanied by clear changes in structure and function of neurons in medial prefrontal cortex (mPFC)^5-7^.Furthermore, mPFC plays a critical role in many forms of decision-making including drug seeking and extinction^8, 9^, thus highlighting the importance of this region for expression of pathological drinking. It is not clear how the computational properties of mPFC are altered during addiction, but alcohol (and other appetitive) cues increase neuronal activity in human medial PFC areas^10, 11^, where activity is linked to perceived reward value^12, 13^. In addition, mPFC plays a key role in guiding decision-making during conflict^14, 15^, and compulsive drinking has been conceptualized as impaired processing of conflict between desire to avoid punishment and drive to consume alcohol^16^.

In animal models, compulsive drinking is frequently operationally defined as aversion-resistant drinking (ARD)^1, 2^. Rodents with heritable risk for excessive drinking rapidly progress to ARD^17, 18^, suggesting a pathophysiology in circuits required to control compulsive drinking. In support of this view, previous work in rats bred for excessive drinking, alcohol preferring rats (P rats), identified alterations in mPFC neural activity that reflected a weakened representation of the “intention” to drink alcohol despite continued consumption^19^, indicating that the role of this brain region in decision-making was compromised.

Outbred rats exhibit compulsive drinking following prolonged alcohol exposure^1,2^, which is mediated by connections from mPFC to ventral striatum^20^. Also, decreased drive from mPFC to periaqueductal gray contributes to compulsive drinking in mice^21^. In heavy-drinking humans, compulsive responding for future alcohol at the risk of immediate punishment corresponded with increased activity in mPFC during alcohol-associated cues^22^. These mPFC findings synergize with behavioral evidence that compulsive drinking involves a “head down and push” strategy^16^ where animals endure the aversive stimulus to obtain alcohol. Therefore, it is critical to determine how altered computational properties of mPFC neural ensembles contribute to expression of compulsive drinking.

To directly examine the importance of mPFC networks for compulsive drinking, we recorded single unit activity from large ensembles of dorsal mPFC (dmPFC, primarily prelimbic and cingulate cortex^23, 24^) neurons in rodent models of compulsive drinking (P rats) and non-compulsive drinking (short drinking-history Wistar rats) during a simple cued-access protocol task. In some drinking sessions, compulsive (ARD) intake was modeled using alcohol adulterated with the bitter tastant quinine (aversive stimulus, “challenged drinking”)^1, 2^. We evaluated neuronal representations of various decision-making relevant signals by dmPFC populations during alcohol and alcohol+quinine sessions using principal component analysis^25^ (PCA). To do so, we developed a novel implementation of PCA capable of identifying stable principal components (PCs) across neural populations of varying size. By analyzing populations of neurons at high temporal resolution^26, 27^ it was possible to examine how large-scale patterns of neural activity evolve during the compulsive decision to drink. We deconstructed the decision-making process into multiple stages by comparing different trials and different time points^28^ including pre-trial seeking state, alcohol-availability cues, sipper approach initiation, and drinking. Critically, we also utilized novel applications of change point analyses^29^ to identify changes in animal seeking state and sipper approach likelihood. With these methods, we tested the hypothesis that representations of behavioral control signals are weakened in dmPFC during compulsive drinking. Finally, emerging data in humans suggests that targeting the mPFC^30-32^ could reduce craving and intake in patients diagnosed with a substance use disorder^33, 34^. Thus, we used a chemogenetic approach to stimulate neurons in dmPFC to test the hypothesis that increasing dmPFC activity would reduce compulsive drinking, which would support the view that circuit manipulations that restore dmPFC function are potentially novel treatments for AUD.

## Results

### P Rats Exhibit Compulsive Drinking

The 2-Way Conditioned Access Protocol (2CAP) task was previously developed by our group to assess cued drinking with or without aversive consequences^19, 35, 36^. Following initial alcohol exposure in home cage (Intermittent Access Protocol (IAP), Fig. 1a), P rats and Wistar rats (Wistars) were trained in the 2CAP task (Fig. 1b). Each 2CAP session contained 48 positive conditioned stimulus (CS+) and 48 negative conditioned stimulus (CS-) trials and was conducted in a chamber with two stimulus lights and retractable sippers on opposite ends of the chamber (Fig. 1c,d). On CS+ trials, a single light (blinking or solid, counter balanced across subjects) was on for 4 seconds and indicated the side of the chamber where 10% ethanol fluid would become accessible for 8 seconds. On CS-trials, both lights were illuminated (opposite modality from CS+) and no access was provided.

**Figure 1:**
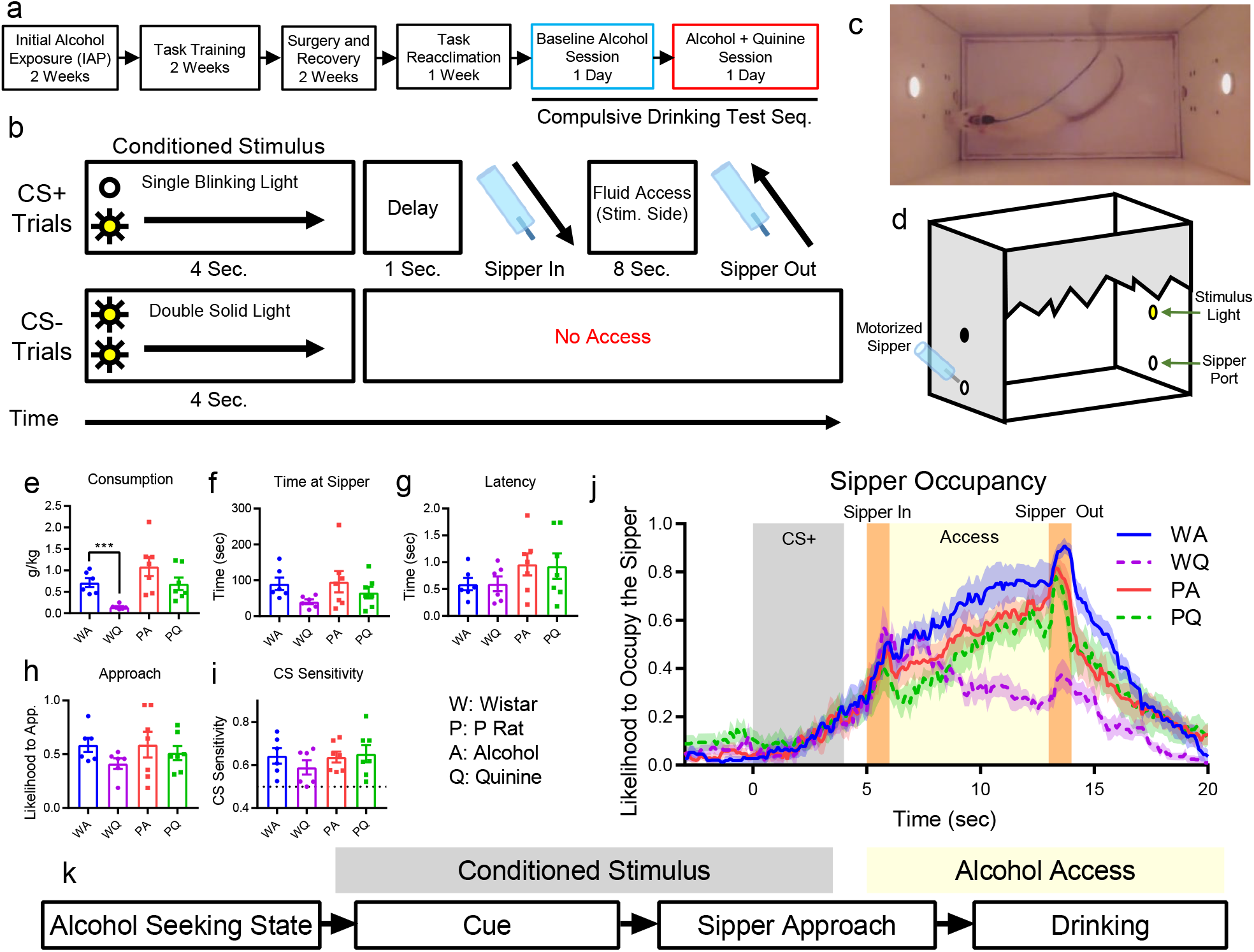
P rats exhibit compulsive drinking. **a**, Experimental schedule for all animals. Electrophysiological data were gathered in two test sessions: 10% alcohol (blue, Tues.) and 10% alcohol + 0.1 g/L quinine (red, Wed.). **b**, 2-Way Conditioned Access Protocol (2CAP) task. On CS+ trials, a stimulus light cued the animal to the side of the chamber where fluid would become available. On CS-trials, both stimulus lights were illuminated and modality of the light (blinking vs. solid) was switched. **c**, Example image of rat in the chamber with both stimulus lights illuminated. **d**, Chamber diagram. **e**, Alcohol consumption dropped significantly for Wistars drinking alcohol + quinine (WQ) vs alcohol-only (WA), but not for P rats (PQ vs PA) (^***^: p<10^−3^). Time at sipper (**f**), latency to approach the sipper (**g**), and likelihood to approach throughout the session (**h**) did not differ with the addition of quinine. **i**, Both Wistars and P rats successfully discriminated between CS+ and CS-(CS Sensitivity = CS+ approaches / all approaches). **j**, Likelihood to occupy the sipper (i.e., to be located at the sipper, in the position to drink, attending the sipper) at a given time during approach trials was similar for all groups up to the beginning of fluid access, where Wistars tended to leave the sipper drinking alcohol + quinine. **k**, We examined dmPFC representations of four key decision-making variables during different task stages: alcohol seeking state (prior to CS onset), alcohol cue (early CS), sipper approach (later CS), and current drinking (access).

Following task training with 10% ethanol (v/v) only, animals were implanted with multielectrode probes (Cambridge Neurotech) in dmPFC. After recovery and task reacclimation, animals underwent a two-day compulsive drinking test sequence that consisted of a baseline regular 2CAP session with 10% ethanol only and a compulsive 2CAP session with 10% ethanol adulterated with 0.1 g/L quinine (Fig. 1a). Electrophysiological data were recorded (OpenEphys) and spike sorted (Kilosort2^37^) for these test sessions.

Replicating our previous work^17^, we found that Wistars decreased consumption significantly when alcohol was adulterated with quinine (Fig. 1e, strain main effect: F(1,22)=9.2, p=0.0062, liquid main effect: F(1,22)=10.05, p=0.0044). Conversely, P rats showed a smaller reduction in consumption of alcohol-quinine, indicating that P rats exhibited compulsive drinking to a greater degree than Wistars. P rats exhibited a preference for unadulterated alcohol during a separate alcohol vs. alcohol+quinine choice test (Supplemental Fig. 1) and it has been shown previously that exposure to quinine at a similar concentration produced elevated aversive facial responses in P rats^38^. These data indicate that P rats could taste the quinine and found it aversive. Based on these results, we will refer to P rats as compulsive subjects and Wistars as non-compulsive subjects in subsequent analyses. Though a trend was observed for Wistars spending less time at the sipper on quinine days (Fig. 1f) and approaching the sipper more quickly (Fig. 1g), no significant strain or alcohol-condition differences were found. Also, no differences were observed in the proportion of trials where the animal approached a sipper (Fig. 1h). All animals were more likely to approach the sipper on CS+ trials relative to CS-trials, indicating that they could differentiate CS+ from CS-(Fig. 1i). As there was no punishment for approaching on a CS-trial, animals still frequently approached on CS-trials. The reduced consumption and time at sipper in Wistars were driven behaviorally by a tendency in alcohol-quinine sessions to approach the sipper, but then leave soon after arriving (Fig. 1j).

### A Novel Method to Assess Neural Population Representations

To precisely define how dmPFC representation of key decision-making variables related to compulsive drinking, in particular why P rats compulsively decided to consume alcohol but Wistars did not, we examined neural representations in dmPFC in four key epochs of the task (Fig. 1k): a pre-trial period to assess neural representation of the animal’s alcohol-seeking state (i.e., seeking or not-seeking), an early CS period to assess neural representation of the alcohol cue, a late CS period to assess neural representation of sipper approach initiation, and an access period to assess neural representation of current drinking. Novel analyses methods were developed to assess behavior and neural activity during these epochs.

We used a new method for applying principal component analysis^25^ (PCA) to examine the representations^39^ of decision-making variables by populations of neurons in dmPFC (Fig. 2a, see Methods for complete analysis work flow). Throughout the analysis, we compared averaged firing rates across pairs of trial types defined either by environmental stimuli (e.g., CS+ vs. CS-) or behaviorally (e.g., seeking vs. not-seeking, drinking vs. not drinking) to assess neural population representations of a given variable that differed across the trial types. Note that we do not attempt to identify which trial represents the signal, rather we only quantify separation as a measure of representation strength. Also, we note that separation between PC trajectories is not necessarily a measure of the subjective strength of a given signal to the animal (i.e., increased PC separation for seeking behaviors does not necessarily imply increased desire for alcohol by the animal).

**Figure 2:**
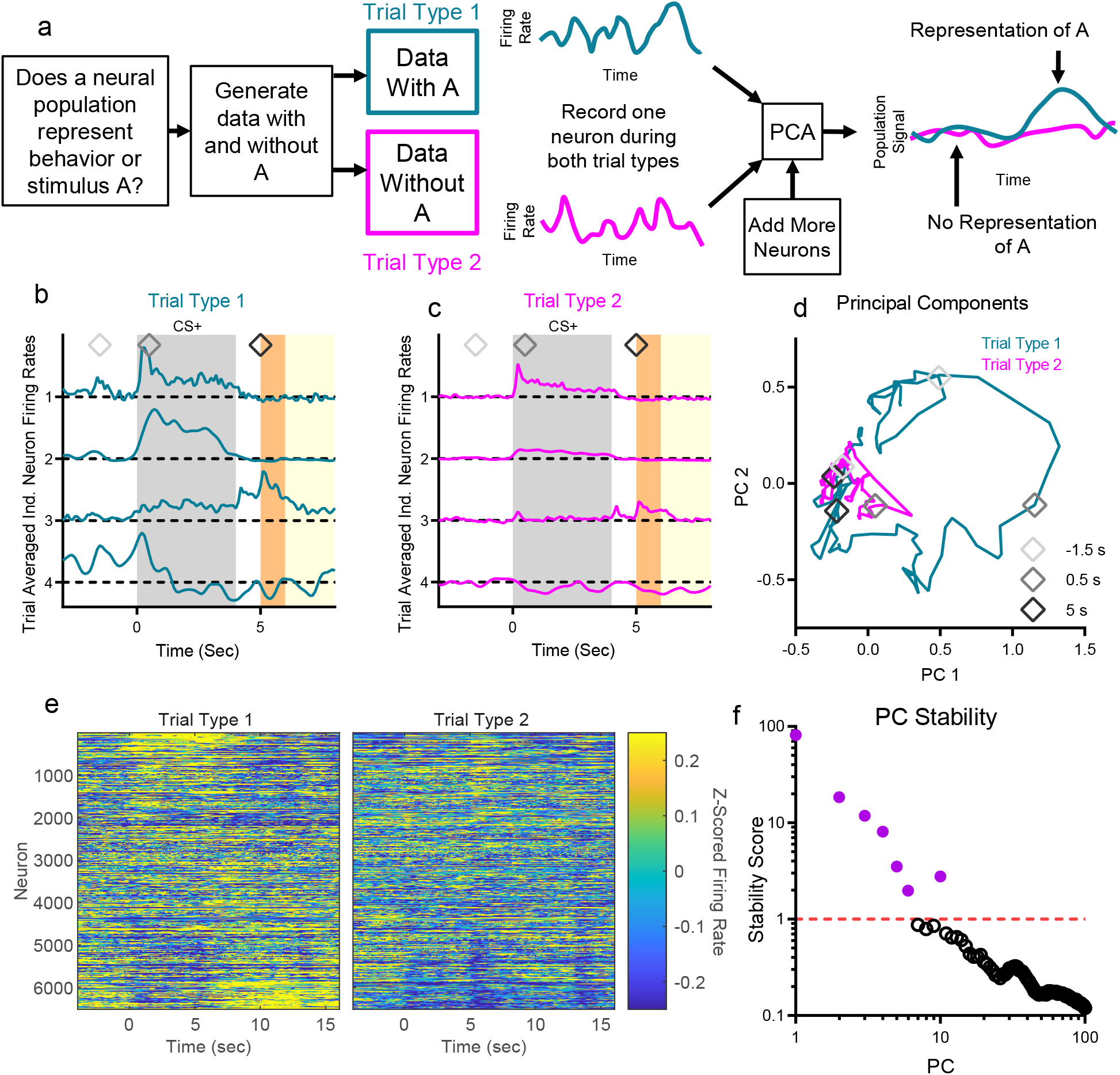
Neural population analysis using principal component analysis. **a**, We used principal component analysis (PCA) to assess neural representations of various decision-making relevant signals by comparing pairs of trial types that differentiate key stimuli (e.g., CS+ vs. CS-) or behaviors (e.g., seeking vs. not seeking, drinking vs. not drinking). **b**,**c**, Four example neurons modulated their trial averaged firing rates dependent on trial type (seeking vs. not-seeking state, see Fig. 3 for more information). **d**, PCA reduced the four neuron firing rates to population signals (PCs) that are linear combinations of individual neuron firing patterns. **e**, Neurons sorted by PC linear combination coefficient (neuron identity matched in right and left panels). Firing responses before CS+ onset, during the CS+, and during access are visible (event timings same as b, but time frame longer in e). **f**, Seven PCs were found to be stable across subsampling trials, which were used to control for differences in neuron yield across experimental groups.

Consider four example neurons recorded from dmPFC (Fig. 2b,c). In this toy example, trial type 1 consists of data recorded when the animal was seeking alcohol and trial type 2 consists of data recorded when the animal was not seeking alcohol (details of behavioral identification of seeking discussed below). Neurons 1 and 2 primarily responded to the CS+, though neuron 2 more clearly differentiated between trial type 1 and trial type 2. Neuron 3 responded to the CS+ and at the beginning of access on trial type 1. Neuron 4 differentiated trial types prior to the CS+ onset. PCA performed on these four neurons produced population signals (principal components (PCs)) that are linear combinations of the individual neuron firing patterns. In this example analysis of four neurons, the PC trajectory for trial type 1 is larger, indicating a more robust population signal (Fig. 2d). Distances between PC trajectories for two types of trials quantifies how well the population of neurons represents differences between those trials at a given time. When we ranked all neurons by their contribution to a selected PC, broad differences between trial types become apparent (Fig. 2e).

When PCA is performed on neural populations of different sizes, there is a risk that the most variance explaining dimensions are mostly driven by the larger population, thus resulting in the spaces from the smaller population being warped to fit the larger one. To address this, we developed a novel method for selecting PCs that relies on subsampling neurons to identify stable PCs across different groups of animals (Fig. 2f, see Methods). These selected PCs accounted for 38% of the overall variance in neural firing.

### Alcohol Seeking State Representation is Enhanced in P Rats and During Challenged Drinking

We assessed the neural representation of the seeking state through differences in firing between seeking and not-seeking states prior to CS+ onset (Fig. 3a). This is a critical stage of the decision-making process because it sets the initial conditions of each cued decision. To assess seeking state prior to each trial, we utilized sipper approach as a key indicator of seeking, where approach is determined by movement to a sipper and into a position to lick from that sipper, as assessed by animal tracking. Animals rarely stayed at one sipper between trials or failed to drink if they approached the correct sipper during access, so the act of approaching represented a necessary behavioral step to obtain alcohol or alcohol+quinine, demonstrating that the animal was seeking to drink. Across animals, the trial-by-trial likelihood to approach gradually decreased throughout a session (Fig. 3b). However, individual animals tended to exhibit a sharp change in approach behavior from a high approach to low approach state (Fig. 3c). We calculated the change point in the likelihood to approach across trials for each animal and marked this as the session change point, which represented the shift from seeking state to not-seeking state (Fig. 3d). No differences were observed between strains or liquid type for session change point (Fig. 3d, Supplemental Fig. 2).

**Figure 3:**
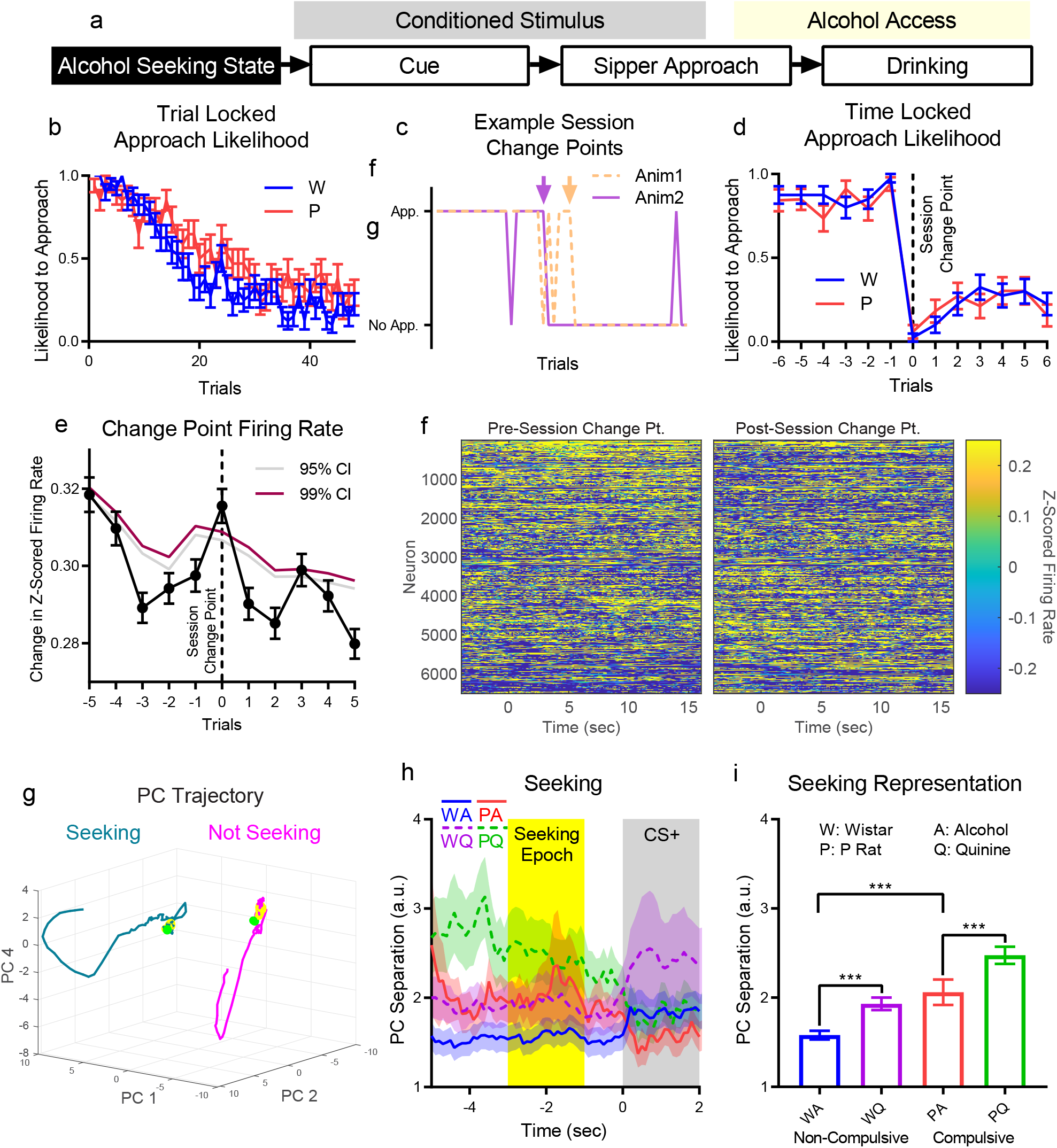
Challenged drinking strengthens seeking representation in dmPFC in compulsive and non-compulsive subjects. **a**, Alcohol seeking was assessed by examining differences in neural population firing patterns prior to task stimuli. When averaged throughout the session, likelihood to approach gradually decreased (**b**), but when time locked to the session change point for each animal, likelihood to approach exhibited a sharp change (**c**: clear examples, arrows mark change point trials, **d**: mean across sessions). Pre- and post-session change point trials were identified as seeking and not-seeking trials, respectively. **e**, Change in average firing rate between trials increased at the session change point above randomized data. (b, d, and e mean+/-sem) **f**, After PCA and sorting neurons by their contribution to PC 1, a large remapping in active neurons was observed across trials immediately before and after the session change point (mean over last/first three trials shown). **g**, Trajectories for seeking and not-seeking states were shifted in population activity space. (Mean trajectory, all neurons. -5 (green dot) to +7 seconds relative to CS+ onset. Yellow sections: seeking epoch (see h).) **h**, PC separation between seeking and not-seeking trials across all stable PCs for each experimental group. (Mean +/-std, only neurons from a given experimental group) **i**, Representation strength increased during challenge (quinine) for both strains and was higher in compulsive subjects. (Mean +/-std shown across mean PC separation values from each 0.1 sec time bin in seeking epoch (h, yellow region). ^***^: p<10^−3^)

Trial by trial change in average firing rate across all neurons exhibited an increase at the session change point above randomized data (Fig. 3e). After performing PCA on neural firing patterns (see Methods) and sorting neurons based on contribution to PC 1, we found a large shift in which neurons were active at the session change point (Fig. 3f). The specific trajectories of the stable PCs throughout the trials were highly varied through the trial, but most PCs possessed large separations before the CS+ in seeking versus not-seeking trials (Supplemental Fig. 3). We focused on a 2-second-long time window (seeking epoch), ending 1 second before CS+ onset, to avoid overlapping in time with CS-signals during intertrial intervals or with the CS+ itself. The effect size for PC separation was calculated for the seeking epoch and PCs 1, 2, and 4 had the largest separation (Supplemental Fig. 4). The trajectories for these PCs were shifted in population space before and after the session change point, reflecting a representation of the seeking state (Fig. 3g). By comparing PC separations across seeking and not-seeking trials for all stable PCs, we could examine differences in the strength of representations of seeking signals through time (Fig. 3h). We examined the mean PC separation for each 100 ms time bin in the seeking epoch and found that compulsive subjects (P rats) possessed stronger baseline seeking representations on alcohol-only days, while challenged drinking (quinine-alcohol) strengthened seeking representations in both strains (Fig. 3i, strain main effect: F(1,38)=429, p<10^−10^, liquid main effect: F(1,38)=237, p<10^−10^, interaction: F(1,38)=27.5, p<10^−5^). Recall that increased seeking representation strength implies neural firing patterns are more distinct in dmPFC, not that animals have an increased desire for alcohol or alcohol+quinine. Collectively, these data show increased seeking representation during challenged drinking and in compulsive subjects during alcohol drinking.

### Cue Representation is Diminished during Compulsive Drinking

The CS+ alerted animals to trial onset and identified the correct sipper where access would be provided. We assessed the neural representation of this cue in two ways (Fig. 4a). First, we compared neural firing patterns shortly before the CS+ onset (−2.4 to -0.5 sec) to neural firing patterns immediately following CS+ onset (0 to 1.9 sec) for all CS+ trials to examine the representation of CS+ onset. PC trajectories for pre-CS+ time windows did not vary for many PCs, consistent with a lack of stimulation or consistent time-locked behavior across trials. In strong contrast, PC trajectories during CS+ showed clear movement through population activity space (Fig. 4b, Supplemental Fig. 5). PC separations rose quickly at CS+ onset and then decayed in all groups (Fig. 4c). When comparing mean PC separation values for each 100 ms time bin in these time windows, challenged drinking increased CS+ onset representation strength in non-compulsive subjects (Wistars), but decreased it in compulsive subjects (P rats) (Fig. 4d, strain main effect: F(1,38)=80.6, p<10^−10^, time main effect: F(19,38)=17.2, p<10^−10^, strain/liquid interaction: F(1,38)=115, p<10^−10^).

**Figure 4:**
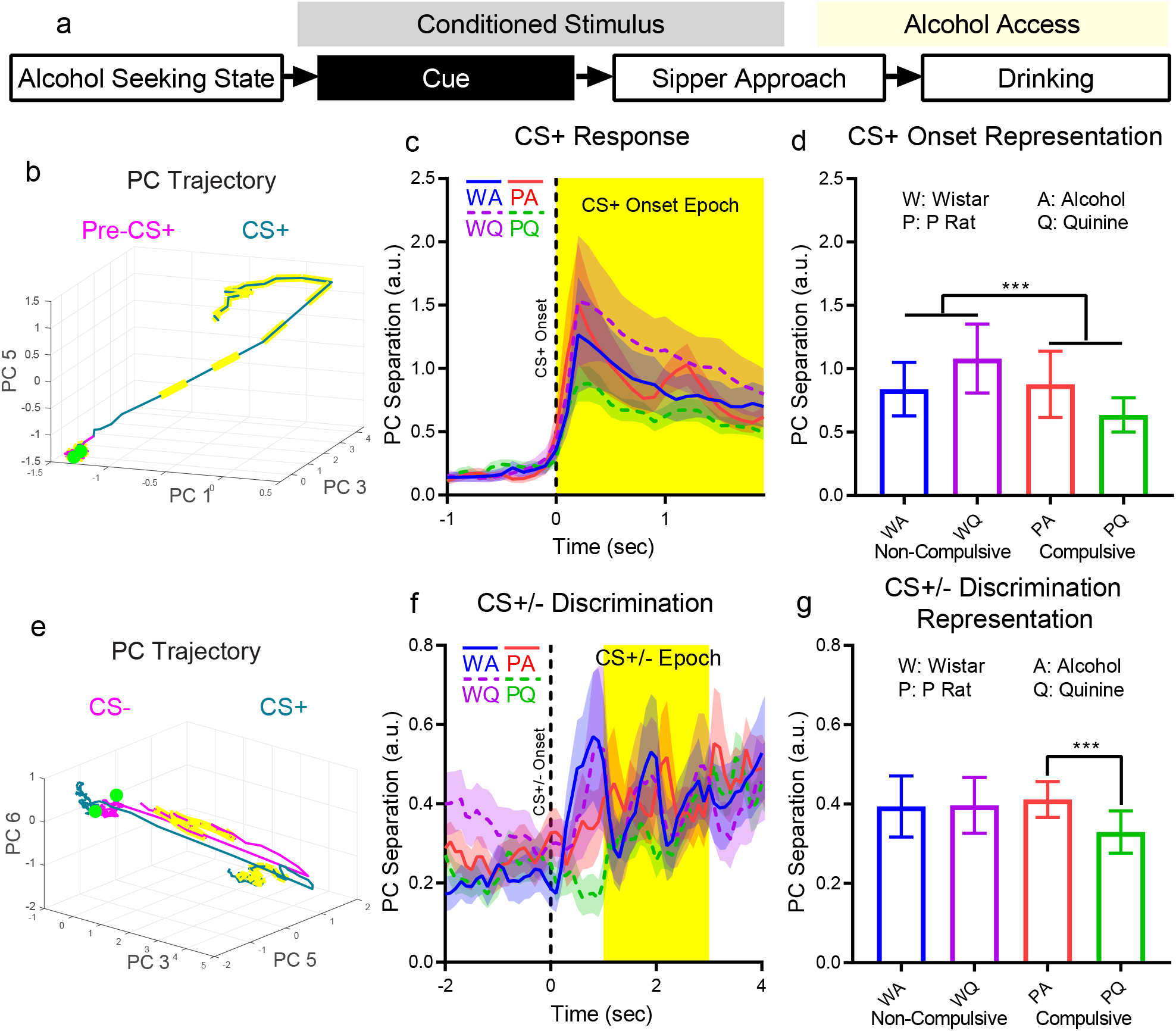
Challenged drinking improved or maintained cue representation in non-compulsive subjects, but weakened cue representation strength in compulsive subjects. **a**, Cue representation was examined by comparing pre-CS+ to CS+ time epochs (CS+ onset representation) and CS+ to CS-time epochs (CS+/-discrimination representation). **b**, Trajectories prior to CS+ onset were stationary, whereas CS+ onset produced a moving trajectory. (−1 (green dot) to +1.9 seconds relative to CS+ onset. Yellow dashed sections: CS+ onset epoch (see c).) **c**, PC separation between pre-CS+ and CS+ time epochs. **d**, Mean PC separation across the CS+ onset epoch (c, yellow epoch) showed that representation strength increased with challenged drinking (quinine) for non-compulsive subjects, but decreased for compulsive subjects. **e**, Trajectories on both CS+ and CS-trials moved quickly from the pre-CS location near the start of the CS, but CS-trials quickly returned towards their original position. (−2 (green dot) to +4 seconds relative to CS+ onset. Yellow sections: CS+/-epoch (see f).) **f**, PC separation between CS+ and CS-trials. **g**, Mean PC separation during the CS +/-epoch (f, yellow time window) showed that CS+/-representation strength was unchanged in non-compulsive subjects during challenge, but decreased in compulsive subjects. ((b) and (e): mean trajectory, all neurons. (c) and (f): mean +/-std, only neurons from a given experimental group, all stable PCs. (d) and (g): Mean +/-std across mean PC separation values from each 0.1 sec time bin in CS+ onset or CS+/-epoch, ^***^: p<10^−3^.)

Next, we compared firing patterns during the CS+ and CS- to examine representations of CS+/CS-discrimination. PC trajectories for CS+ and CS-trials were similar following the initial CS onset, but CS-trajectories returned to the pre-CS position in activity space while CS+ trajectories did not (Fig. 4e, Supplemental Fig. 6). All groups exhibited increased PC separation during the CS+/-epoch (Fig. 4f). [Oscillations seemed to correspond to the on/off cycle of the blinking/solid light modality used for the CS+/- that was counter-balanced across animals.] CS+/-representation strength (across the CS epoch) decreased only for compulsive subjects (P rats) in challenged drinking (Fig. 4g, liquid main effect: F(1,38)=8.70, p=0.0054, strain/liquid interaction: F(1,38)=9.93, p=0.0032). Overall, cue representation strength was similar in compulsive and non-compulsive subjects when drinking alcohol, but diverged under challenge such that strength in non-compulsive subjects increased or stayed the same, but strength decreased in compulsive subjects.

### Approach Initiation Representation is Enhanced in Non-Compulsive Subjects but Diminished in Compulsive Subjects During Challenged Drinking

Sipper approach initiation is the first physical action in the consumption process. We focused on neural activity the end of the CS to examine this signal (Fig. 5a). We utilized a novel method to determine initiation of sipper approach. We calculated the instantaneous likelihood to approach the sipper at every time bin after CS+ onset using the animal’s position and velocity (Fig. 5b, see Methods). On approach trials, the change point in this likelihood to approach marked the time point where the animal initiated approach (Fig. 5c). On no-approach trials, random approach initiation times relative to CS+ onset were used for time locking with approach trials. When we time locked neural firing patterns to the approach-initiation time point, we found a wide range of dynamic response patterns near the approach-initiation time point (Fig. 5d-f, Supplemental Fig. 7). For instance, PCs 3 and 10 showed widening separations between approach and no-approach trials starting approximately 5 seconds before approach initiation. PC 4 showed very small separations prior to approach initiation, at which point the separation between approach and no-approach trials grew. Though movement differences were found between approach and no-approach trials (Supplemental Fig. 8), these differences were stable through time prior to approach initiation, and correlations between PCs and animal speed were low (Supplemental Fig. 9), indicating that these PC results are not merely movement artifacts, though they may represent motor planning. PC trajectories for no-approach trials were relatively compact, whereas PC trajectories for approach trials showed movement through population activity space from an initial point several seconds before approach initiation (Fig. 5g). Despite the variety of dynamic patterns during approach initiation, the total PC separation for each experimental group was relatively stable through time (Fig. 5h). To assess the approach-initiation representation, a time window was selected -3 to -1 seconds before approach initiation (Fig. 5h, yellow window) to reduce any effects of movement initiation near the approach-initiation time point. The mean PC separation for each 100 ms time bin in this epoch was higher in non-compulsive than compulsive subjects during alcohol-only drinking (similar to our previous work^19^), and increased in non-compulsive subjects, but decreased in compulsive subjects during challenge drinking (Fig. 5i, strain main effect: F(1,38)=1332, p<10^−10^, liquid main effect: F(1,38)=27.0, p<10^−5^, strain/liquid interaction: F(1,38)=202, p<10^−10^). Therefore, approach initiation strength was depressed in compulsive subjects drinking alcohol, while challenged drinking increased representation strength in non-compulsive subjects and depressed it yet further in compulsive subjects.

**Figure 5:**
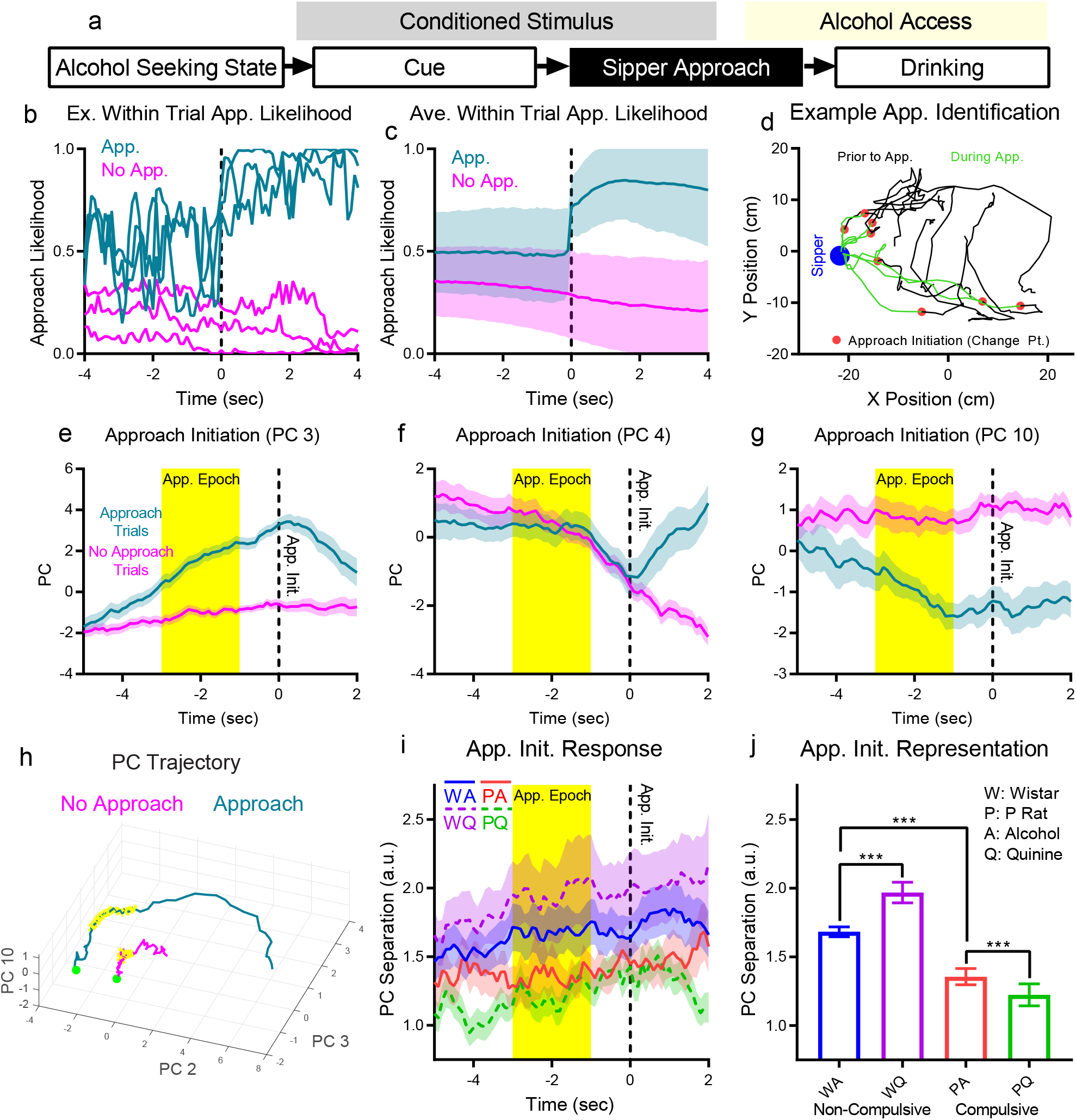
Challenged drinking strengthened the representation of the sipper approach initiation signal in non-compulsive subjects, but weakened the representation in compulsive subjects. **a**, The representation of sipper approach initiation was examined by comparing approach to no-approach trials just before approach initiation. Change points in instantaneous likelihood to approach the sipper in each trial (**b**: examples, **c**: averages) were used to identify sipper approach initiation. **d**, Eight example approach trials from an example animal that clearly show the range of approach initiation behaviors captured by this method. **e-g**, Approach and no-approach trials began to diverge several seconds before approach initiation for PCs 3 and 10. Conversely, PC 4 only showed a divergence following approach initiation, indicating that this PC may track movement. (All PCs shown in Supplemental Fig. 8.) **h**, Large differences in PC trajectories were observed between approach and no-approach trials. (Mean trajectory, all neurons. -5 (green dot) to +2 seconds relative to approach initiation. Yellow sections: approach epoch (see h).) **i**, PC separation between approach and no-approach trials all stable PCs. (Mean +/-std, only neurons from a given experimental group.) **j**, Mean PC separation across the pre-approach initiation epoch (h, yellow time window) showed that the representation was stronger in non-compulsive subjects on alcohol days^19^ and that representation strength increased with challenge (quinine) in non-compulsive subjects, but decreased for compulsive subjects. (Mean +/-std shown across mean PC separation values from each 0.1 sec time bin in (f). ^***^: p<10^−3^)

### Drink Representation is Enhanced in Non-Compulsive Subjects During Challenged Drinking

The final signal we examined was the neural population representation of drinking by comparing CS+ drink and CS+ no drink trials (Fig. 6a). Drink initiation times were identified by tracking data and randomly selected drink initiation times relative to CS+ onset time were used as time lock points for no drink trials. PC trajectories for no-drink trials were relatively compact while trajectories for drink trials showed large divergences in population activity space (Fig. 6b, Supplemental Fig. 10). PC separations (during the 2 seconds following drinking initiation) for non-compulsive subjects during challenged drinking possessed a large peak immediately following the onset of drinking (Fig. 6c), indicating that dmPFC firing patterns in non-compulsive subjects diverged substantially more when drinking alcohol+quinine when compared to other experimental groups, indicating a stronger representation (Fig. 6d, strain main effect: F(1,38)=155, p<10^−10^, liquid main effect: F(1,38)=92, p<10^−10^, strain/liquid interaction: F(1,38)=133, p<10^−10^). This drink representation increase may underlie Wistar rejection of quinine-alcohol, while the lack of drink representation change may allow continued compulsive drinking in P rats.

**Figure 6:**
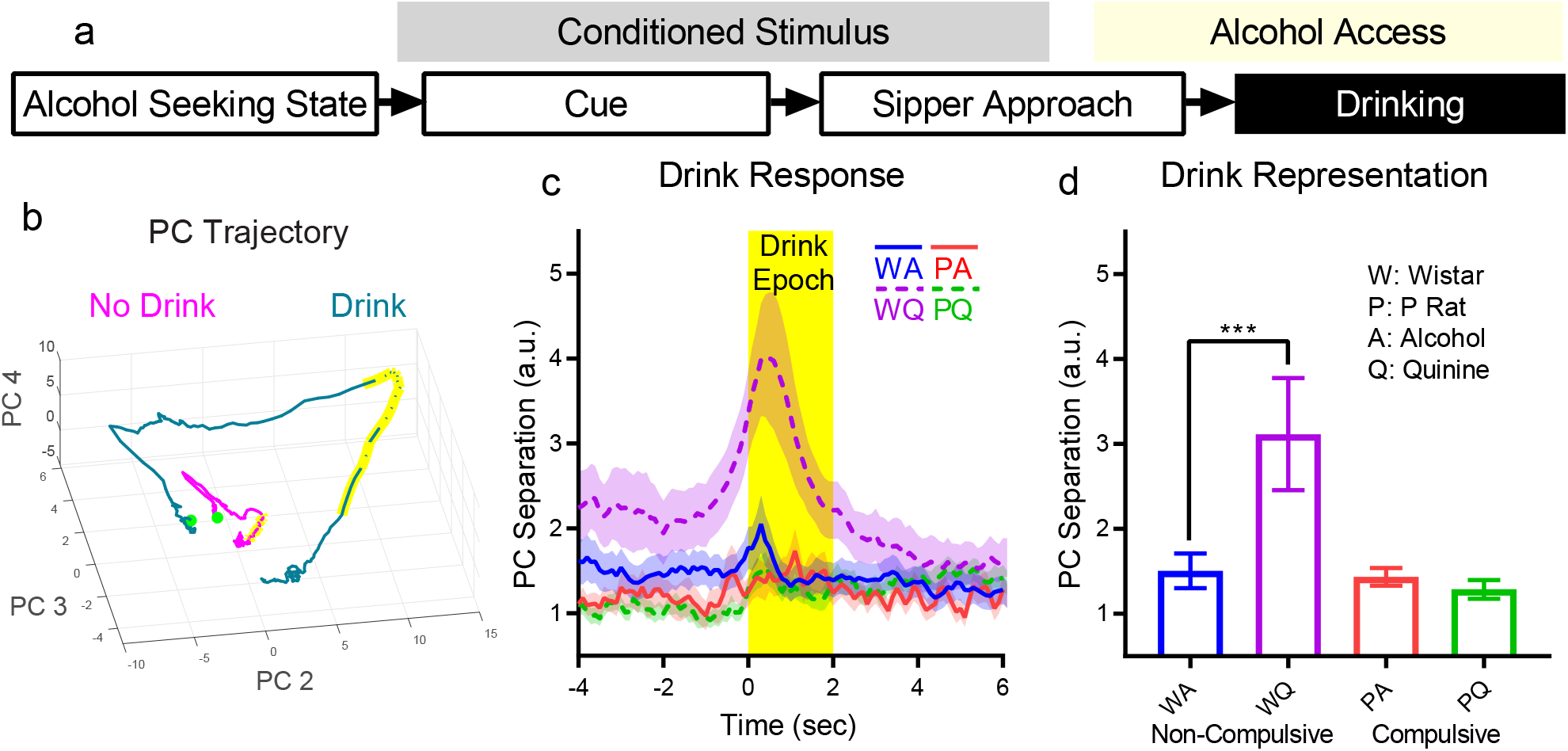
Challenged drinking strengthened dmPFC representation of drinking in non-compulsive subjects, but not in compulsive subjects. **a**, The representation of drinking was assessed by comparing neural firing patterns around the initiation of drinking, specifically with regards to differences between trials where the animal drank and trials where it did not. No drink trials used randomly selected drink initiation delays relative to CS+ onset. **b**, Drink trial trajectories showed a large loop in population activity space, whereas no drink trials were more compact. (Mean trajectory, all neurons. -10 (green dot) to +10 seconds relative to drink initiation. Yellow sections: drink epoch (see c).) **c**, PC separation across all stable PCs during times immediately before and during drinking onset (Mean +/-std, only neurons from a given experimental group). Non-compulsive subjects during challenged drinking (quinine) showed a large peak in PC separation indicating a large change in population activity in response to quinine that was absent in other groups (i.e., a stronger representation of drinking). **d**, Mean PC separation in the drink epoch (c, yellow time window) showed strengthened representations during challenged drinking (quinine) in non-compulsive subjects. (Mean +/-std shown across mean PC separation values from each 0.1 s time bin in drink epoch. ^***^: p<10^−3^)

### dmPFC Excitation in P Rats Prevents the Progression of Compulsive Drinking

To better understand the large divergence in firing patterns in non-compulsive subjects during challenged drinking (Fig. 6d), which coincided with marked reductions in alcohol consumption (Fig. 1j), we increased the data set with additional challenged drinking test sessions, and examined drink representation and individual neuron firing responses during the drink epoch (Fig. 7a-f). Furthermore, we analyzed neuron waveforms to differentiate excitatory and inhibitory neurons^40, 41^ (Supplemental Fig. 11). In non-compulsive subjects, challenged drinking continued to produce elevated drink representation and suppressed intake in later compulsive-drinking tests (Fig. 7a, Supplemental Fig. 12, test session main effect: F(1,38)=13.5, p=0.00073, liquid main effect: F(1,38)=244, p<10^−10^). The increase in drink representation during challenge (quinine-alcohol) corresponded to increased firing in both excitatory (liquid main effect: F(1,3448)=18.8, p<10^−4^) and inhibitory (liquid main effect: F(1,210)=7.27, p=0.0076) neurons (Fig. 7b,c). However, in compulsive subjects, drink representation increased during additional drinking sessions regardless of liquid type and intake remained elevated (Fig. 7d, Supplemental Fig. 12, test session main effect: F(1,38)=161, p<10^−10^, liquid main effect: F(1,38)=21.7, p<10^−4^, time main effect: F(19,38)=2.09, p=0.026, test session/liquid interaction: F(1,38)=10.5, p=0.0024), which corresponded to increased firing in excitatory neurons only (Fig. 7f, test session main effect: F(1,2523)=66.88, p<10^−10^). (See Supplemental Fig. 13 for inhibitory neuron firing time series.) Two later recordings (one Wistar and one P rat) were switched between compulsive and non-compulsive groups due to reversed compulsive drinking behavior (Supplemental Fig. 14).

**Figure 7:**
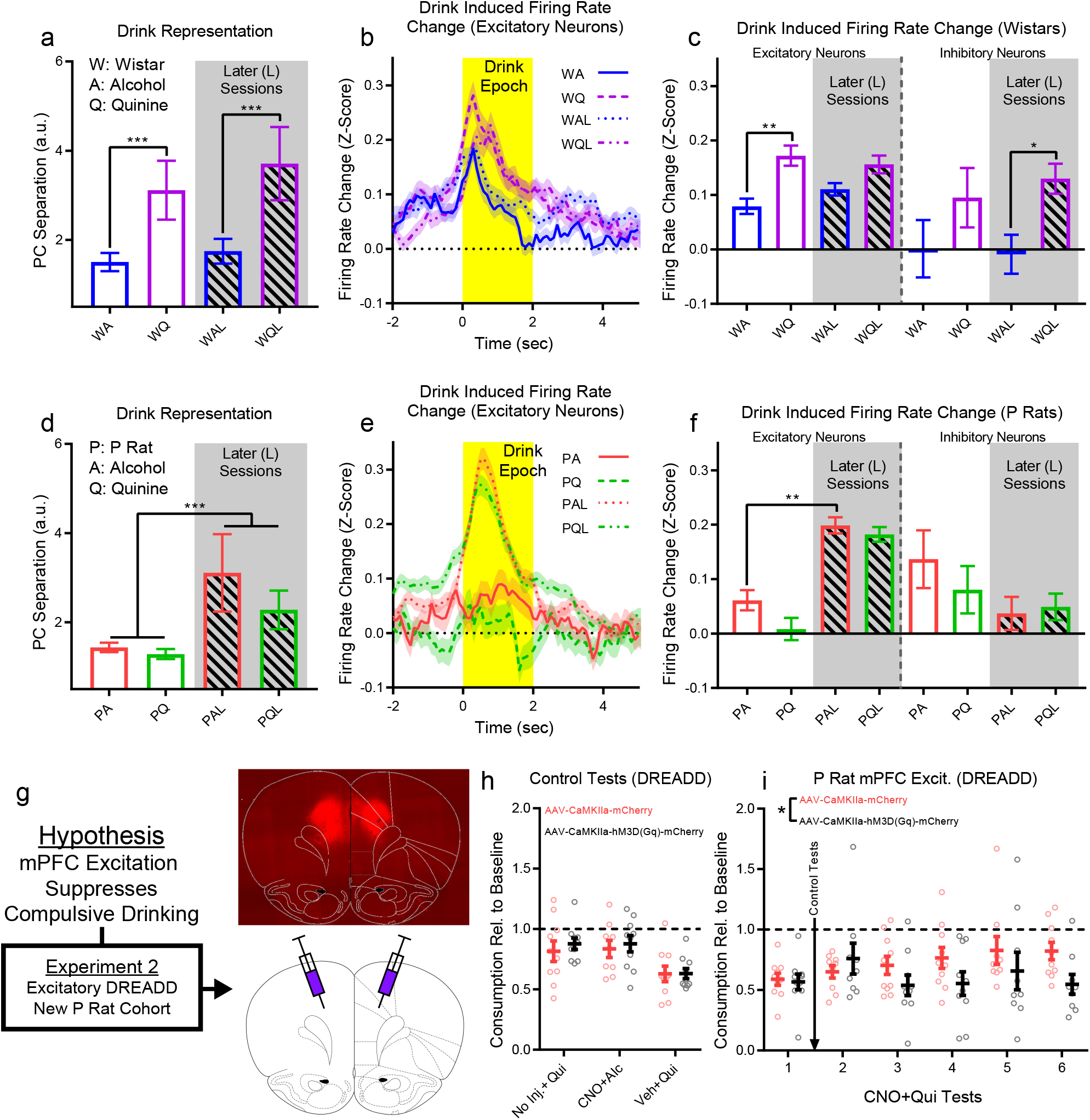
Changes in drink representation are related to excitatory/inhibitory neuron behavior and DREADD-mediated dmPFC excitation prevented the progression of compulsive drinking. **a**, Increased drinking representation strength in non-compulsive subjects was persistent in later challenged drinking tests. This corresponded to increased excitatory (**b, c**) and inhibitory (c, Supplemental Fig. 13) neuron firing. **d**, Compulsive subjects developed increased drinking representation during both challenged and unchallenged drinking, corresponding to increased firing in only excitatory neurons (**e, f**). **g**, Thus, non-compulsive animals exhibited an increase in excitatory neuron firing on quinine-alcohol days relative to alcohol-only that was *not* observed in compulsive animals. Therefore, we hypothesized that driving dmPFC excitatory neurons would suppress compulsive drinking in compulsive animals. To test this hypothesis, a new cohort of 20 P rats underwent bilateral excitatory CAMKII-a DREADD or control virus injections in dmPFC (exemplary expression image). **h**, Animals expressed compulsive drinking, alcohol consumption was only slightly reduced by CNO, but consumption decreased for challenged drinking during vehicle injection, regardless of virus expressed. **i**, After multiple challenged drinking tests, control animals exhibited increased compulsive drinking, while CNO/DREADD remained relative unchanged. (a and d, mean +/-std shown across mean PC separation values from each 0.1 s time bin in drink epoch (first session results reproduced for comparison (Fig. 6d)). b and e, mean +/-sem change in firing rate on drink trials. c and f, mean +/-sem of mean neuron firing rate change in drinking epoch. For all graphs, ^***^: p<10^−3, **^: p<10^−2^, *: p<0.05)

Since increased firing during challenged drinking was associated with reduced intake in Wistar rats, we hypothesized that dmPFC excitation suppresses compulsive drinking. Therefore, we conducted a second experiment in a new cohort of 20 P rats using identical alcohol exposure, 2CAP training, and compulsive drinking test procedures (Fig. 7g). We injected Designer Receptors Exclusively Activated by Designer Drugs (DREADDs: AAV5-CaMKIIa-hM3D(Gq)-mCherry, control: AAV5-CaMKIIa-mCherry) in dmPFC in this cohort of animals. These animals exhibited only a small reduction in drinking during challenged drinking with quinine (compulsive drinking) and no injection, as well as when drinking alcohol during DREADD-mediated increased excitation following a CNO injection (3 mg/kg) (Fig. 7h). However, injection of vehicle (i.e., no DREADD-mediated increase in excitation) paired with challenged drinking decreased consumption in both DREADD and control animals (test main effect: F(2,53)=8.09, p<0.001), though not as substantially as was observed in Wistars (Fig. 1e). During challenged drinking with CNO injections, we did not find a difference between animals with DREADD-mediated increased excitation and controls during the first test (Fig. 7i). However, repeated tests showed that consumption of alcohol+quinine remained constant in DREADD-mediated increased excitation animals while it increased in control animals (Fig. 7i, Supplemental Fig. 15, DREADD main effect: F(1,103)=5.34, p=0.023). CNO/DREADD did not alter locomotion (Supplemental Fig. 15). Thus, dmPFC excitation over time prevented the progression of compulsive drinking.

## Discussion

Overall, we propose that challenged intake in compulsive drinkers was driven by a strong seeking representation and weak representations of behavioral control variables, such as cues, approach initiation, and drinking. We labelled these as “behavioral control” variables because they represented the immediate, trial-by-trial signals that determined consumption on a given trial and because they changed in similar ways across groups. The seeking variable was different in that it operated on a longer time scale (i.e., across multiple trials). While the seeking variable was inferred behaviorally by actions governed by behavioral control variables (i.e., high likelihood to approach implied the animal was seeking), we conceptualized the seeking state of the animal as antecedent to behavioral control.

When grouped in this way, challenged drinking in non-compulsive subjects resulted in increased engagement in both seeking state and behavioral control representations (Fig. 8). Conversely, challenged drinking in compulsive subjects resulted in a shift toward an enhanced seeking state representation and away from choice specific variable representations. (Note that, while drinking representation increased during alcohol+quinine sessions in compulsive subjects for later exposures, it was still similar to or lower than the drinking representation during later alcohol sessions.) Therefore, it may be possible to prevent compulsive drinking by increasing the representation of behavioral control variables in dmPFC during challenged drinking.

**Figure 8:**
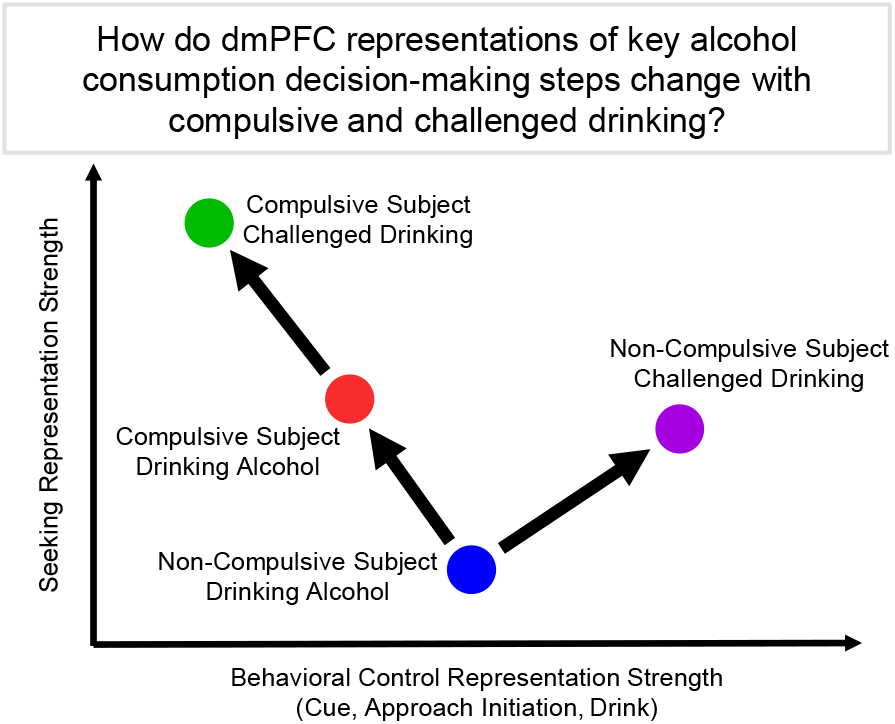
Main conclusions. Using electrophysiology in compulsive drinking (P rats) and non-compulsive drinking (Wistars) subjects during their first challenged drinking test, we examined representations in dmPFC of seeking state and three behavioral control variables: cue, approach, and drinking. We found that dmPFC in non-compulsive subjects more robustly represented behavioral control and seeking state signals when drinking was challenged by an aversive stimulus. For compulsive subjects, relative to non-compulsive subjects, dmPFC better represented alcohol seeking state, but represented a key behavioral control variable worse when drinking alcohol. When drinking was challenged in compulsive subjects, the representation of behavioral control signals weakened and the representations of seeking strengthened yet further in dmPFC

A striking result of these studies was the large increase in drink representation immediately after drinking onset in non-compulsive subjects (Wistars) during challenged drinking. Furthermore, in these subjects we found that excitatory and inhibitory neurons tended to increase their firing rate immediately after drink onset. This increase in drink representation strength and individual neuron firing response persisted through multiple compulsive drinking tests in non-compulsive subjects. Conversely, compulsive subjects (P rats) exhibited no change in drink representation or individual neuron firing rate during the first compulsive drinking test. During later compulsive drinking tests, compulsive subjects exhibited a large increase in drink representation and excitatory neuron firing during both unchallenged and challenged drinking sessions.

While these patterns of drink representation increases are similar, there are subtle differences in the conditions that produce these increases that point to possible explanations for dmPFC’s role in compulsive drinking. Non-compulsive subjects exhibited increases in drink representation only during challenged drinking, when their expectation about the contents of the solution was violated, leading to response inhibition and decreased consumption. This behavior is similar to previous reports that the anterior cingulate cortex (ACC, a region overlapping dmPFC^24^) responds to unexpected negative outcomes (so called error-related negativities)^42^ and that these large scale signals are caused by altered neural activity across many neurons^43^. Furthermore, neurons in ACC have been shown to respond to conflict during response inhibition in stop-go or stop-change tasks^44, 45^. Also, it has been argued that increased excitation and inhibition in close temporal proximity – as we observed in only non-compulsive subjects during challenged drinking – improves network performance and encoding^46^. Therefore, we hypothesize that increased drink representation during challenged drinking in non-compulsive subjects was the neural signature of conflict processing in dmPFC that contributed to decreased consumption. Compulsive subjects exhibited increased drink representation during both challenged and unchallenged drinking in later sessions. A possible explanation for increased drink representation in later unchallenged drinking session is that the initial quinine challenge subsequently altered the expectation of the contentions of the reinforcer, whereby the possibility of receiving unadulterated alcohol became less certain. We suggest that the increased drink representation during later challenged sessions relative to earlier sessions is a result of malfunctioning conflict processing that results in a head-down-and-push behavioral strategy^16^. Importantly, non-compulsive animals exhibited an increase in the drink representation during challenged drinking relative to unchallenged drinking, which was absent in compulsive animals in the first and later sessions. The absence of this representation increase in compulsive subjects could be due to many factors, including changes within dmPFC, changes in upstream brain regions that encode rewarding and aversive stimuli, and/or the connections from those upstream regions to dmPFC. Identifying the precise neural circuits that are responsible for this diminished representation during compulsive drinking is a crucial question for future studies.

Based on the increase in excitation in non-compulsive subjects during challenged drinking that we observed and transcranial magnetic stimulation studies in human substance use disorder patients^33, 34^, we hypothesized that exciting dmPFC in P rats would reduce compulsive drinking. To test this hypothesis, we expressed excitatory DREADDs in pyramidal neurons (CAMKIIa promoter) in dmPFC in P rats. We found that activation of the DREADD did not eliminate but prevented the escalation of compulsive drinking over subsequent challenged drinking sessions. Therefore, we hypothesize that excitation of dmPFC pyramidal neurons disrupted maladaptive processes necessary for the escalation of compulsive drinking.

The decision to consume is a particularly important facet of the three-stage cycle theory of addiction (binge/intoxication, withdrawal/negative affect, preoccupation/anticipation)^47, 48^. This decision point is the critical intermediary between the preoccupation/anticipation stage and the binge/intoxication stage and therefore plays a key role in perpetuating this cycle. This study demonstrated differences between compulsive and non-compulsive subjects in a brain region established to play a critical role in decision-making – the dmPFC. These data indicate that, during the decision to consume, compulsive drinking subjects exhibit enhanced seeking and diminished behavioral control representations in dmPFC. This difference in compulsive drinking subjects may facilitate consumption despite negative consequences, thereby continuing the addiction cycle by moving the subjects to the binge/intoxication stage.

## Online Methods

### Subjects

This study was conducted using male alcohol preferring (P) rats (Indiana University School of Medicine) and male Wistar rats (Envigo, Indianapolis). All animals were shipped via ground transportation from breeding facilities to the laboratory, all within the city of Indianapolis. All animals were given food and water ad libitum in their home cages. All animal procedures were approved by the Indiana University Animal Care and Use Committee.

The electrophysiology study utilized 24 P rats and 72 Wistars over 4 sub-cohorts. Following IAP and 2CAP training, 8 P rats and 8 Wistars were selected for surgery (two sub-cohorts of 4 Wistars and two sub-cohorts of 4 P rats). One P rat and one Wistar were excluded from these sub-cohorts due to damaged probes, resulting in a final yield of 7 P rats and 7 Wistars. Electrophysiological data for one Wistar was lost for its first ARD test, so it was excluded from the analysis of the first ARD test. Later ARD test data were intact, so they were included in those analyses.

The DREADD experiment utilized 23 P rats. 20 animals were selected for virus surgery following IAP and 2CAP training (10 with DREADD, 10 Control). One DREADD animal was lost prior to the third from last test session.

### Intermittent Access Protocol (IAP)

An intermittent access protocol (IAP) was used to acclimate animals to the taste and effects of alcohol^35, 49^. 20% ethanol (Decon Laboratories, Inc.) in one bottle and tap water in another bottle was provided ad libitum in the animals’ home cages for 24 hour periods on alternating days. Animals were weighed and bottles were placed on the cages on Monday, Wednesday, and Friday mornings (approximately 2 hours into the animals’ dark cycle) for 2 weeks. Bottles were pulled 24 hours later and weighed to assess consumption. Animals were selected for 2CAP task training based on their average consumption across IAP (Supplemental Fig. 16). Within each sub-cohort of animals, we selected lower drinking P rats (with the exception of four P rats that did not drink) and higher drinking Wistars in an attempt to bring group consumption means closer together. We also removed animals with health concerns.

### Two-Way Conditioned Access Protocol Task (2CAP)

The Two-Way Conditioned Access Protocol (2CAP)^19, 35, 36^ task was used to assess voluntary consumption of 10% ethanol in tap water. All training sessions and DREADD testing sessions were conducted in standard rat shuttle chambers (MedAssociates) housed in custom sound attenuating boxes. Regular 2CAP sessions and baseline ARD testing sessions used 10% ethanol (Decon Laboratories, Inc.) in tap water. Quinine test sessions used a 10% ethanol solution adulterated with 0.1 g/L quinine (Fisher Scientific). Electrophysiological data were gathered in a custom made chamber to allow for sufficient space above the animal’s head for the headstage and wiring. We utilized an updated version of the 2CAP task that included negative conditioned stimuli (CS-). Two possible stimuli modalities were used: a solid 4 second light on stimulus or a 4 second stimulus composed of four 300 ms light on followed by 700 ms light off blinks. Positive conditioned stimuli consisted of one modality of stimulus presented directly above the sipper where fluid would become available. Negative conditioned stimuli consisted of the other stimulus modality presented above both sippers. The specific modality type assignment (e.g., solid light CS+, blinking light CS-) was counter balanced across subjects. The pseudorandom nature of the CS+ and CS-trials resulted in a slightly elevated likelihood (approximately 60% instead of 50%) for animals to receive a CS-trial following a CS+ trial and vice-versa. Each 2CAP session lasted approximately 1 hour and consisted of 48 CS+ and 48 CS-trials. CS+ trials were separated by pseudorandomly selected intertrial intervals of 20, 28, 36, 44, 56, 68, 96, and 120 seconds. CS-trials randomly occurred within the intertrial intervals, though an exclusion period of 3 seconds at the start and end of CS+ trials prevented CS+ and CS-trials for occurring adjacently. Animals had a low likelihood to approach both sippers (i.e., approach the correct side after approaching the incorrect side) or approach the sipper prior to the end of the CS+ (Supplemental Fig. 17).

Animals were selected for surgery based on their average consumption across 2CAP training sessions (Supplemental Fig. 16). Within each sub-cohort of animals, we selected lower drinking P rats and higher drinking Wistars in an attempt to bring group consumption means closer together. We also removed animals with health concerns. Blood ethanol concentration levels were correlated with consumption (Supplemental Fig. 18).

### Electrophysiology

Animals selected for electrophysiological recordings underwent implantation surgery using procedures approved by the Indiana University Animal Care and Use Committee. Animals were anesthetized using isoflurane and held in a stereotax (Kopf). The scalp of the animal was shaved and cleaned with alcohol and betadine. Cefazolin (30 mg/kg) was administered IP and Bupivacaine (5 mg) was injected subcutaneously in the scalp. After removing the scalp and clearing the bone, bregma and lambda were located. Grounding screws were implanted posterior to lambda above the cerebellum and 6 mounting screws were implanted (4 between lambda and bregma, 2 approximately 5 mm anterior of bregma). A trephine was used to perform a craniotomy above medial prefrontal cortex. For all animals except one P rat, a multishank probe (Cambridge NeuroTech, 64 channels on 4 or 6 shanks) was attached to a moveable drive and implanted with target coordinates of 3.2 mm A/P, 0.8 mm M/L, 3.0 mm D/V^23^ in the right hemisphere with a 7.5 degree angle in towards the midline. The probe shanks were aligned so they spanned the anterior/posterior direction. One P rat was implanted with a fixed single shank probe (Cambridge NeuroTech, 32 channels) with target coordinates of 3.2 mm A/P, 0.6 mm M/L, 4.3 mm D/V^23^ in the right hemisphere with a 7.5 degree angle in towards the midline. Movable drives were incased in antibiotic ointment and then affixed to the skull using dental cement. Ketofen (5 mg/kg) was administered and the animal was monitored for 7 days post-surgery.

Following recovery, subjects were given two 2CAP sessions in a specialized electrophysiology chamber to reacclimate to the task and acclimate to the new chamber. Next, subjects underwent at least 3 2CAP sessions with electrophysiology wires attached to acclimate to the presence of the wiring. Omnetics headstages were plugged in to the exposed connectors from the neural probes on the dental cement headcap. Electrophysiological data and task specific variables (e.g., stimuli) from the Med Associates controller were recorded using Open Ephys data acquisition systems. Video of the animal was recorded from above the chamber using a Logitech webcam. Animal tracking was performed offline using DeepLabCut^50^. Based on the manual identification of drinking trials and the tracking information, the drinking position of each animal was identified and the drinking initiation time was set as the first time point when the animal’s snout entered the drinking position (within 9 pixels (< ∼1 cm) of the manually identified sipper location). Spike sorting was performed using Kilosort2^37^. Moveable probes were lowered by approximately 0.1 mm the day before each ARD test baseline session. Typically, the probes were lowered after the subject ran a regular 2CAP session on Monday to prepare for ARD baseline and test days on Tuesday and Wednesday.

Final positions of shanks were verified via histology (Supplemental Fig. 19) and quantified (QuickNII-WHSRat-v3)^51^. There was a slight, but significant difference in mean medial/lateral placement of 0.11 mm between P rats and Wistars. There was a larger, significant difference in mean anterior/posterior placement of 0.81 mm between P rats and Wistars. To ensure that differences in recording sites could not explain differences in neural encoding, the correlation between neuron location (defined as the location of the shank on which the neuron was recorded) and neuron principal component coefficients for each stable principal component (see below) was calculated. Across all combinations of PC and the spatial dimension, the *R*^*2*^values for these correlations were less than 0.0117, with most values less than 0.001. This indicates that these differences in recording location cannot explain the observed differences in neural encoding between P rats and Wistars. Recordings from 6 Wistars in the first ARD test yielded 655 neurons on alcohol days and 697 neurons on alcohol+quinine days. Recordings from 7 P rats in the first ARD test yielded 383 neurons on alcohol days and 304 neurons on alcohol+quinine days. Recordings from 7 Wistars for later ARD tests (14 total tests) yielded 1181 neurons on alcohol days and 1235 neurons on alcohol+quinine days. Recordings from 7 P rats for later ARD tests (10 total tests) yielded 1015 neurons on alcohol days and 1044 neurons on alcohol+quinine days. No correlation was observed between neuron firing rate and amount of alcohol consumed^52^.

Among the later compulsive drinking electrophysiology sessions, two subjects were switched between groups based on alcohol and quinine consumption (see Supplemental Fig. 14).

Neurons were classified as inhibitory or excitatory neurons using their largest amplitude mean waveform (Supplemental Fig. 11). The waveform was rescaled from -1 (maximum hyperpolarization amplitude) to 1 (maximum post-action potential depolarization peak). Similar to previously used methods^40, 41^, we classified the neurons using time delays associated the waveform depolarization. We used the time to 50% depolarization and the time to 95% depolarization as the time delays to characterize the speed of the depolarization. When these delays for each waveform were plotted, two clusters were apparent. Waveforms were manually clustered into inhibitory (fast depolarization), excitatory (slower depolarization), and outliers based on these time delays.

### DREADD Experiments

Animals selected for DREADD experiments underwent virus injection surgery using procedures approved by the Indiana University Animal Care and Use Committee. Animals were anesthetized using isoflurane and held in a stereotax (Kopf). The scalp of the animal was shaved and cleaned with alcohol and betadine. Cefazolin (30 mg/kg) was administered IP and Bupivacaine (5 mg) was injected subcutaneously in the scalp. After removing the scalp and clearing the bone, bregma was located. Bilateral craniotomies were performed above dmPFC. Bilateral injections of 0.65 uL at 0.2 uL/min of virus were performed using Hamilton syringes and Harvard Apparatus pumps using the target coordinates of 3.2 mm A/P, 0.8 mm M/L, 3.0 mm D/V^23^ with a 20 degree angle in towards the midline. Syringes were left in place for 10 minutes following injection and then removed slowly over 5 minutes. Ketofen (5 mg/kg) was administered and the animal was monitored for 7 days post-surgery. Virus expression was verified in all animals by histology (Supplemental Fig. 20).

Animals receiving the excitatory DREADD were given AAV5-CaMKIIa-hM3D(Gq)-mCherry (AddGene) and control animals received AAV5-CaMKIIa-mCherry (UNC Vector Core). Tests in which the DREADD was activated were conducted using 3 mg/kg Clozapine N-oxide (CNO, Cayman Chemical Company Inc.) in 1% dimethylsulfoxide (DMSO) and saline IP injections approximately 30 minutes prior to session start.

DREADD experiments took place in the same Med Associates shuttle chambers used for 2CAP training. Lick data was used to identify errors in consumption measures from bottle leaks using the following procedure. For each animal, number of licks and consumption for all baseline and test sessions were linearly fit. Then, the residuals for each session were calculated. The residuals for all sessions for all animals were then compared and residuals above three standard deviations were excluded. This resulted in 4 out of 197 test measurements being excluded.

### Electrophysiology Data Analysis

Data analysis of spike trains was conducted using the following steps (see Supplemental Fig. 21). The raw spike trains (30 kHz) were rebinned using 100 ms bins (10 Hz). Each spike train was then smoothed using an adaptive Gaussian kernel with a standard deviation of one-quarter of the mean interspike interval for that neuron. We will refer to the smoothed spike train of neuron *ii*at time bin *t* as *x*_*i*_(*t*). Next, based on the four stages of the decision to consume alcohol that were discussed (seeking state, cues, initiating approach, and drinking), we extracted segments of the spike train to form trials of interest.

For seeking state analyses, we focused on pre- and post-session change point CS+ trials. Let *t*_*cs*+,*j*_ represent the time bin *t* of the *j*th CS+ trial relative to CS+ onset. We extracted segments of the spike trains near the CS+ (−10 seconds before CS+ onset to 20 seconds after CS+ onset). Let *a*_*i,j*_(*t*)represent the smoothed spike train of neuron *ii*at time bin *t* ∈ [−100,200] relative to *t*_*cs*+,*j*_ on trial *j*. Next, we z-scored *a*_*i,j*_(*t*)across all time bins and trials. Then, we averaged across trials before the session change point trial (*j*_*scp*_) to obtain a mean high seeking state trail 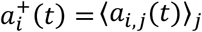 such that 1 *≤ j < j*_*scp*_) and we averaged across trials after the session change point trial to obtain a mean low seeking state trial (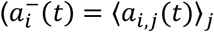 such that *j*_*scp*_ *≤ j ≤ j*_*max*_). The session change point trial was identified by constructing a sequence of approach (both correct and incorrect sipper, coded as 1) and no-approach (coded as 0) in time order for each CS+ trial. This sequence was analyzed using the Matlab function findchangepts.m.

For analyses of drinking cues, we focused on CS+ and CS-trials. Let *t*_*cs,j*_ represent the time bin *t* of the *j* th CS trial relative to CS onset. We extracted segments of the spike trains near the CS (−10 seconds before CS onset to 20 seconds after CS onset). Let *b*_*i,j*_(*t*) represent the smoothed spike train of neuron *i* at time bin *t* ∈ [−100,200] relative to *t*_*cs,j*_ on trial *j* Next, we z-scored *b*_*i,j*_(*t*)across all time bins and trials. Then, we averaged across CS+ trials to obtain a mean CS+ trail 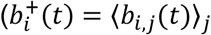 such that *j* is a CS+ trial) and CS-trials to obtain a mean CS-trial (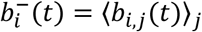 such *j* is a CS-trial).

For approach initiation analyses, we focused on approach (both correct and incorrect sipper) and no-approach CS+ trials. Trials were marked as approach trials if the animal approached either sipper during the access period. The approach initiation time was identified using a change point analysis. Let 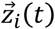 represent the position and velocity of the animal’s snout at time bin *t* relative to the CS+ onset on trial *i*. zUsing both the position and velocity of the animal’s snout allowed the algorithm to account for differences in movements to approach the sipper based on different initial positions and velocities at the start of the trial. Let *p*_*j*_ represent the approach status for trial *j* (1 for approach trials and 0 for no-approach trials). We calculated the likelihood that the animal would approach at time bin *t* on trial *i*(*q*_*i*_(*t*)) using the weighted average of the approach status (*q*_*i*_(*t*)*= ∑*_*j≠i*_*w*_*i,j*_ (*t*)*p*_*j*_) where the weights were the normalized inverse Euclidean distance between the locations and velocities of the animal’s snout at the same time bin relative to CS+ onset on two trials (i.e., 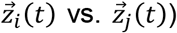). This sequence was analyzed using the Matlab function findchangepts.m to identify the change point in the approach likelihood which represents the approach initiation time bin on trial *i*(*t*_*AI,i*_). For no-approach trials, the approach initiation time relative to the CS+ onset was randomly selected from the approach initiation times relative to the CS+ onset from approach trials for that animal on that day. We extracted segments of the spike trains near the approach initiation time (−5 seconds before approach initiation to 2 seconds after approach initiation). Let *c*_*i,j*_(*t*)represent the smoothed spike train of neuron *i* at time bin *t* ∈ [−50,20]relative to *t*_*AI,j*_ on trial *j*. Next, we z-scored *c*_*i,j*_(*t*)across all time bins and trials. Then, we averaged across approach trials to obtain a mean approach trail (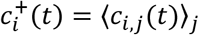such that *j* is an approach trial) and no-approach trials to obtain a mean no-approach trial (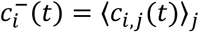 such that *j* is a no-approach trial).

For the analysis of drinking, we focused on correct approach trials and not correct approach trials (i.e., incorrect approach and no-approach trials). We observed that animals that correctly approached (i.e., were in the drinking position at the extended sipper during access) always licked the sipper at least once, though we were unable to resolve the precise lick time. Therefore, we refer to correct approach trials as drink trials. Let *t*_*D,j*_ represent the time bin of the *j*th CS+ trial when the animal first arrived at the sipper (i.e., drink time). For no drink trials (i.e., trials where the animal did not approach either sipper or approached the incorrect sipper), we randomly selected drink times relative to the CS+ onset from drink times relative to the CS+ onset on drink trials. We extracted segments of the spike trains near the drink time (−10 seconds before drink time to 10 seconds after drink time). Let *d*_*i,j*_(*t*)represent the smoothed spike train of neuron *ii*at time bin *t* ∈ [−100,100] relative to *t*_*D,j*_ on trial *j*. Next, we z-scored *d*_*i,j*_(*t*)across all time bins and trials. Then, we averaged across drink trials to obtain a mean drink trail (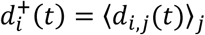 such that *j* is a drink trial) and no drink trials to obtain a mean no drink trial (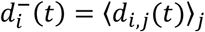 such *j* is a no drink trial).

All trial means of interest were concatenated through time for each neuron (i.e., the sequence 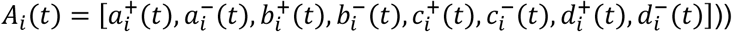) and principal component analysis (PCA) was performed on 500 iterations of 200 randomly selected neurons from each group (animal strain, liquid type (alcohol vs. alcohol+quinine), and ARD test number (first test vs. later tests)). We ran the PCA with neurons considered as variables and time bins considered as observations in order to capture patterns of neural activity across neurons at each time bin of the trials of interest. This process of randomly subsampling neurons and matching the number of neurons from each experimental group in the PCA prevented one group from dominating the PCA, but allowed the PCA to be meaningful across groups.

For each principal component (PC), we determined the stability of the PC across the 500 iterations using the following method. Let e_*i,j*_(*t*)represent principal component *ii*at time bin *t* in the concatenated sequence of trials on PCA iteration *j*. PCs that were found to be inverses of the mean PC across all iterations were inverted by multiplication with -1. For each PC, we calculated the ratio of the variance in the mean PC throughout time to the variance across iterations using *var*_*t*_*⟨*e_*i,j*_(*t*)*⟩*_*j*_/*var*_*t*_e_*i,j*_(*t*)− *⟨*e_*i,j*_(*t*)*⟩*_*j*_. If the variance ratio was above 1, we considered the PC to be stable across PCA iterations and used it in subsequent analyses.

PCs for each experimental group (e.g., Wistars drinking alcohol+quinine in the first compulsive drinking test) were generated by projecting from only the 200 neurons from that group that were used in each of the 500 iterations. Let *f*_*i,j,k*_ represent the coefficient for neuron *ii*for PC *j* on PCA iteration *k*. The projection for PC *j* on PCA iteration *k* for a given experimental group were calculated as *E*_*j,k*_ (*t*)*= ∑*_*i∈G*_ *f*_*i,j,k*_ *A*_*i*_(*t*) such that *G* is the set of 200 neurons from the experimental group of interest that were randomly selected for this PCA iteration. Unless otherwise stated, the mean across 500 iterations was calculated for PC projections by experimental groups and for all neurons together. PC separations at each time bin were calculated as the Euclidean distance between all stable mean PCs for the two trials types of interest (e.g., high seeking vs. low seeking) at corresponding time points in the trials (e.g., 2 seconds before CS+ onset).

A leave one out analysis was performed by rerunning the analysis with every combination of one animal removed. Only minor qualitative differences were observed. Furthermore, a randomized analysis was performed by shuffling assignments for the trial types of interest (e.g., before and after session change point when assessing seeking state). PC stability and PC separations were substantially reduced, indicating that the signals observed herein represent true neural population signals associated with the decision-making signals of interest.

### Statistics

For comparisons between P rats and Wistars across alcohol sessions and alcohol+quinine sessions, we utilized a two-way ANOVA to detect main effects of rat strain and liquid type, as well as an interaction. We used the same procedures for other two-way comparisons, such as first compulsive drinking test sessions vs. later sessions and alcohol vs. alcohol+quinine sessions. For comparisons between P rats and Wistars across alcohol and alcohol+quinine sessions that also incorporated measurements through time (e.g., comparisons of PC separations during epochs of interests), we utilized a three-way ANOVA to detect main effects of rat strain, liquid type, and time, as well as strain*liquid and strain*liquid*time interactions. In the main text, we report only significant (p<0.05) results of these comparisons. We followed these ANOVAs with post-hoc Bonferroni corrected t-Tests (behavior, Fig. 1) or Tukey-Kramer tests (all other figures) and we report the results for three comparisons of interest: Wistars/alcohol vs. Wistars/alcohol+quinine, P Rats/alcohol vs. P Rats/alcohol+quinine, and Wistars/alcohol vs. P Rats/alcohol. Significant results of these post-hoc comparisons are reported in figures only. These post-hoc comparisons were not performed or reported in the case of a significant main effect of time in three-way ANOVAs and relevant other significant main effects are marked in figures instead. We used the same two-way approaches for the analyses of DREADD vs. control animals and multiple drinking sessions.

Throughout figures, error bars and fringe represent standard error of the mean (sem), with the exception of PC separation plots and their summary bar graphs where we show standard deviation (std). We chose to show standard deviation for these graphs because the sample size was controlled by the number of subsamples used in the PCA method in those cases. In all figures, we identify in the caption the type of method used for each error bar or fringe to avoid confusion.

### Data and Software

All spike sorted data, behavioral data, and analysis software are supplied as a supplemental.

## Supplemental

**Supplemental Figure 1:**
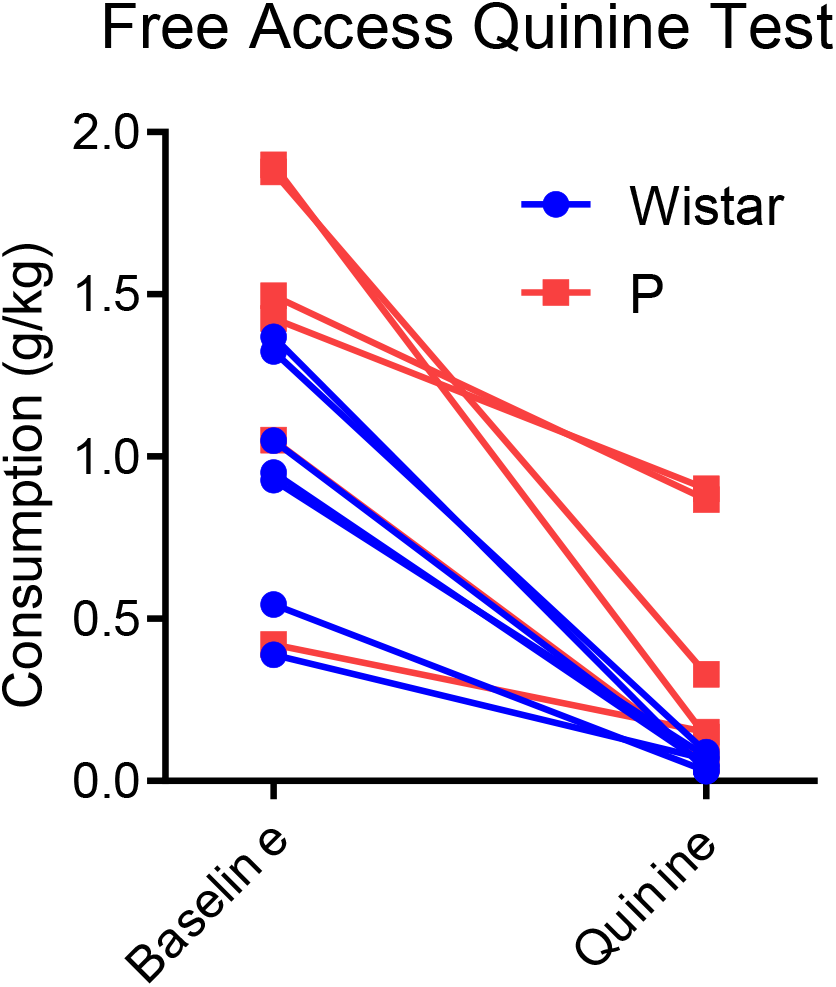
Free access quinine test. Animals were given approximately 60 minutes of free access to both sippers filled with 10% ethanol on baseline days (total time matched to regular 2CAP session, sippers extended into the chamber throughout the whole session). During a subsequent session, the preferred sipper from the baseline session was adulterated with 0.1 g/L quinine. During this quinine session, the animal was manually required to sample both sippers twice. For one P rat, the non-preferred sipper was accidentally adulterated with quinine for which we show quinine consumption from the non-preferred sipper. This animal consumed more quinine adulterated alcohol than regular alcohol during quinine testing. Another P rat was lost prior to this testing. All other animals demonstrated a preference for regular alcohol over alcohol adulterated with 0.1 g/L quinine.

**Supplemental Figure 2:**
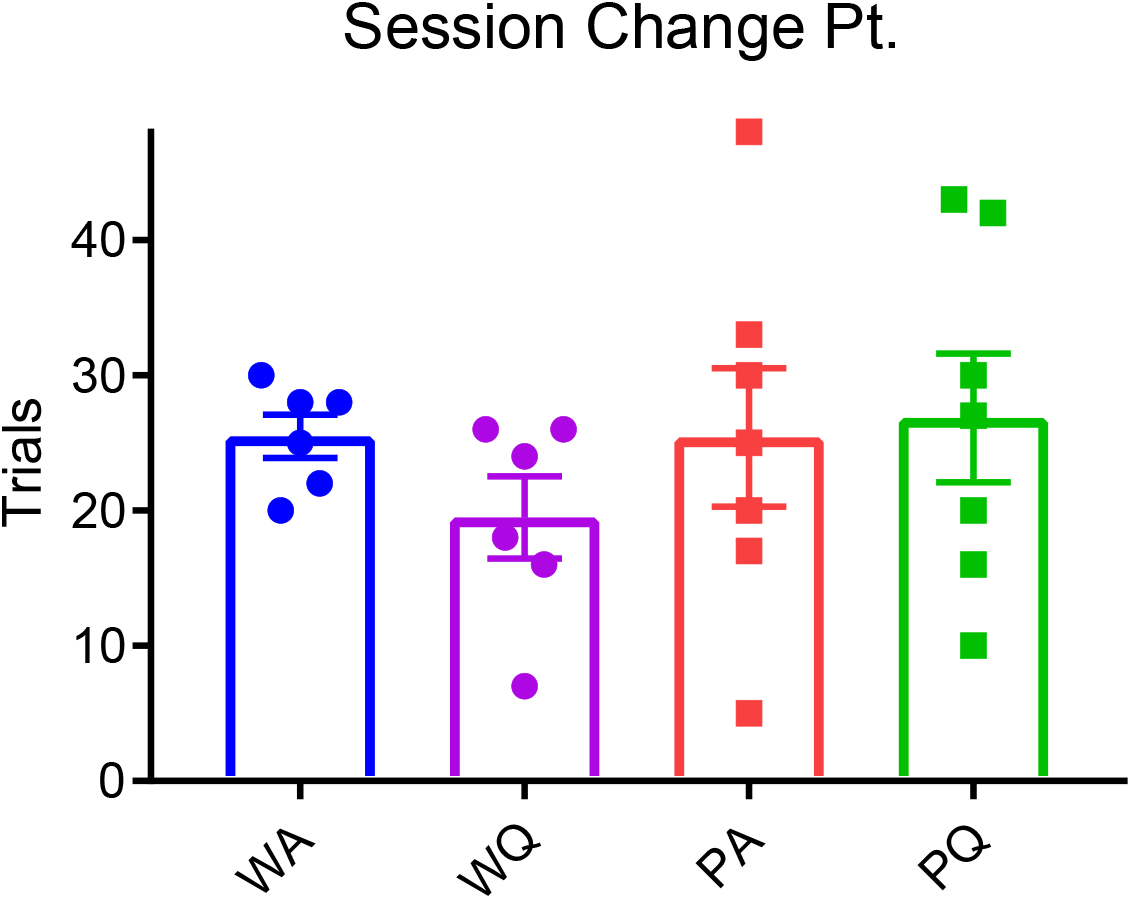
Session change point was not affected by animal strain or liquid type.

**Supplemental Figure 3:**
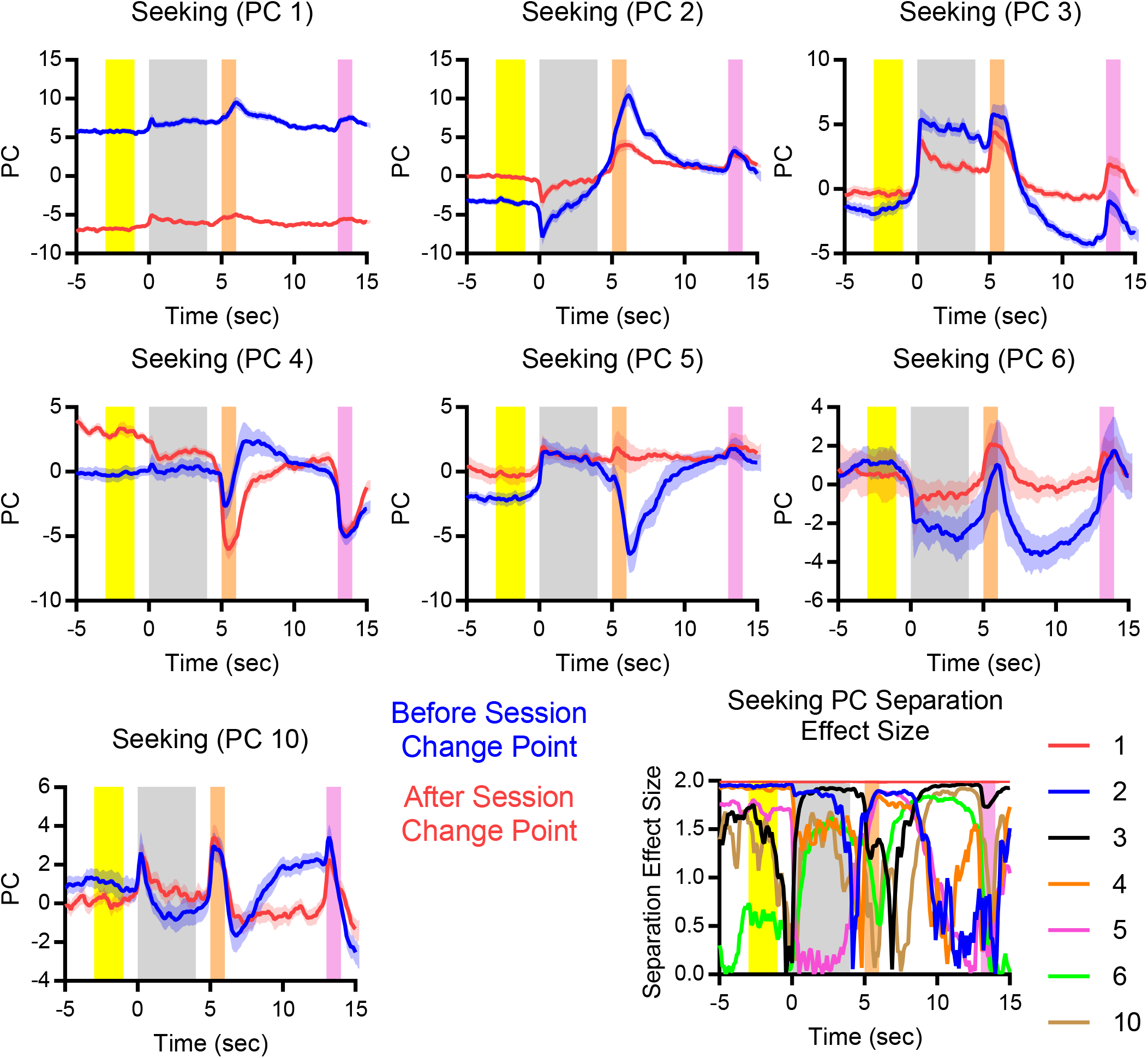
All stable PCs (all neurons) for trials before and after session change point (mean +/-std). Recall, approaching decreased substantially following session change point, marking a change in seeking state. Seeking representation was assessed during the yellow epoch prior to CS+ onset. Many PCs (especially 1, 2, and 4) showed robust separation during the seeking epoch.

**Supplemental Figure 4:**
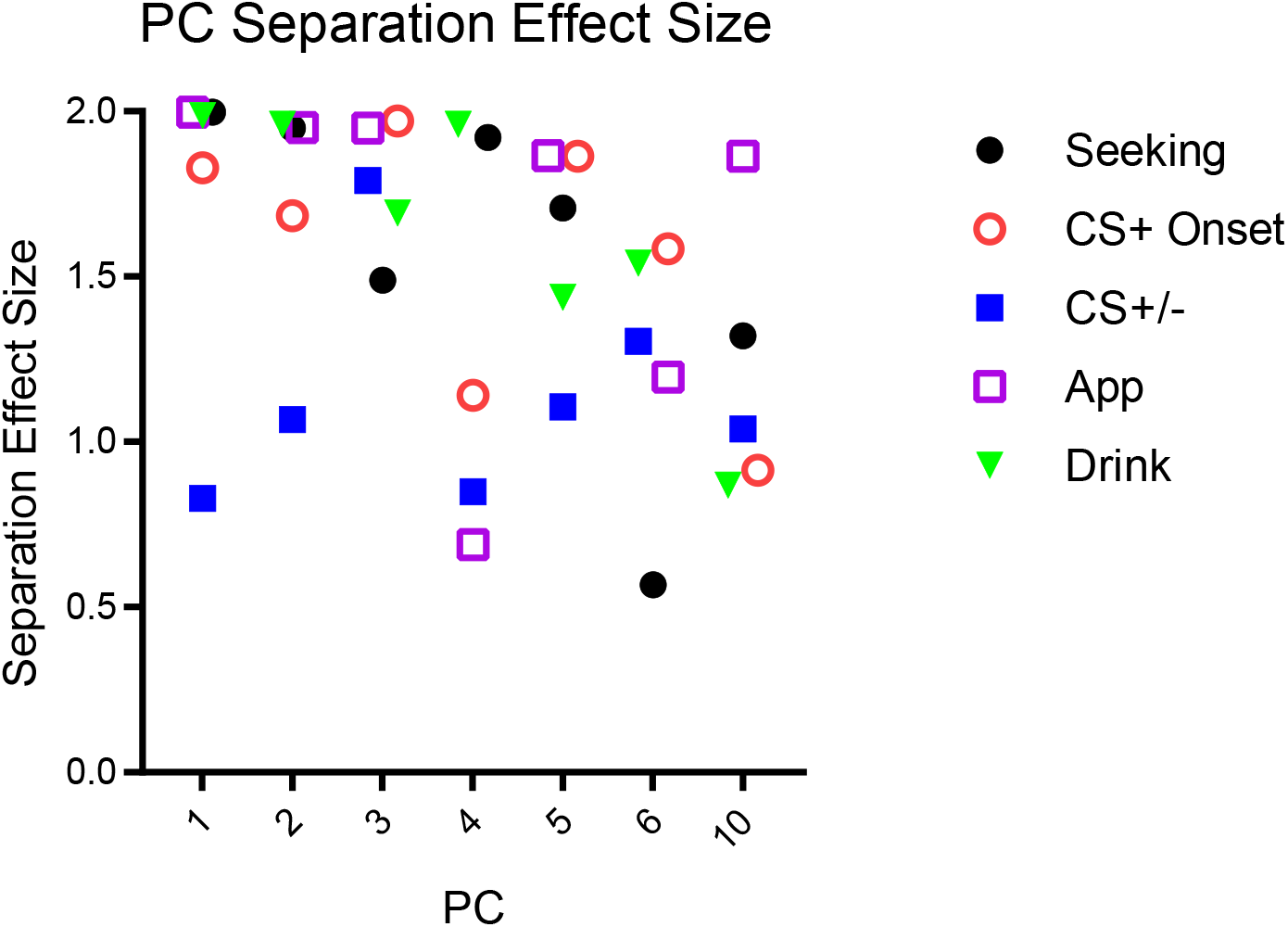
PC separation effect sizes across epochs of interest for each type of representation. Effect size was calculated using Cohen’s d using individual PCA subsampling trials and the epochs of interest.

**Supplemental Figure 5:**
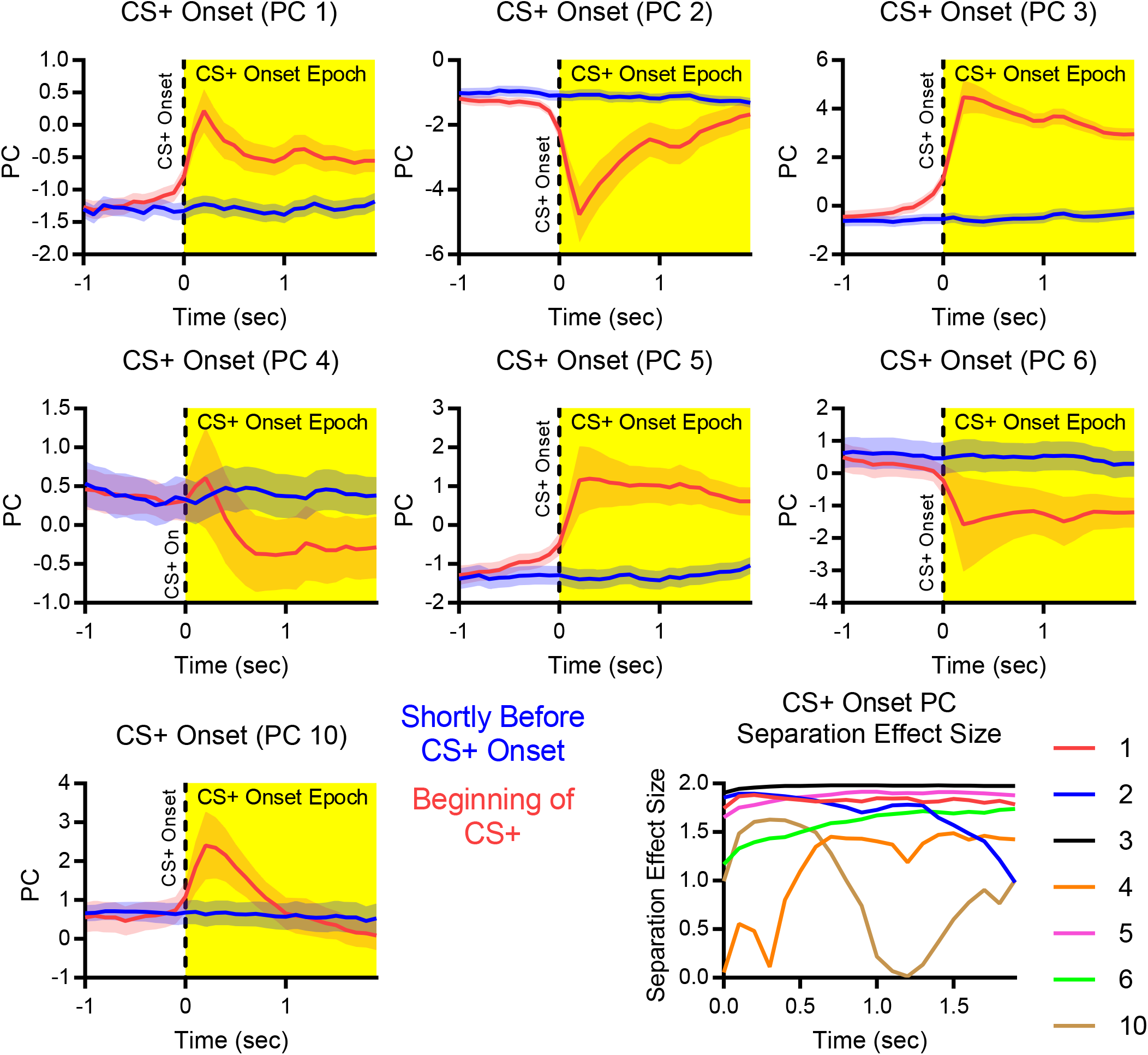
All stable PCs (all neurons) averaged over 2 seconds shortly before CS+ onset (−2.4 to -0.5 seconds before CS+ onset) and during the first 2 seconds of the CS+ (0 to 1.9 seconds after CS+ onset) (mean +/-std). CS+ onset representations were assessed by comparing these epochs, though more time shown for clarity. Many PCs (especially 3 and 5) showed robust separation during the CS+ onset epoch.

**Supplemental Figure 6:**
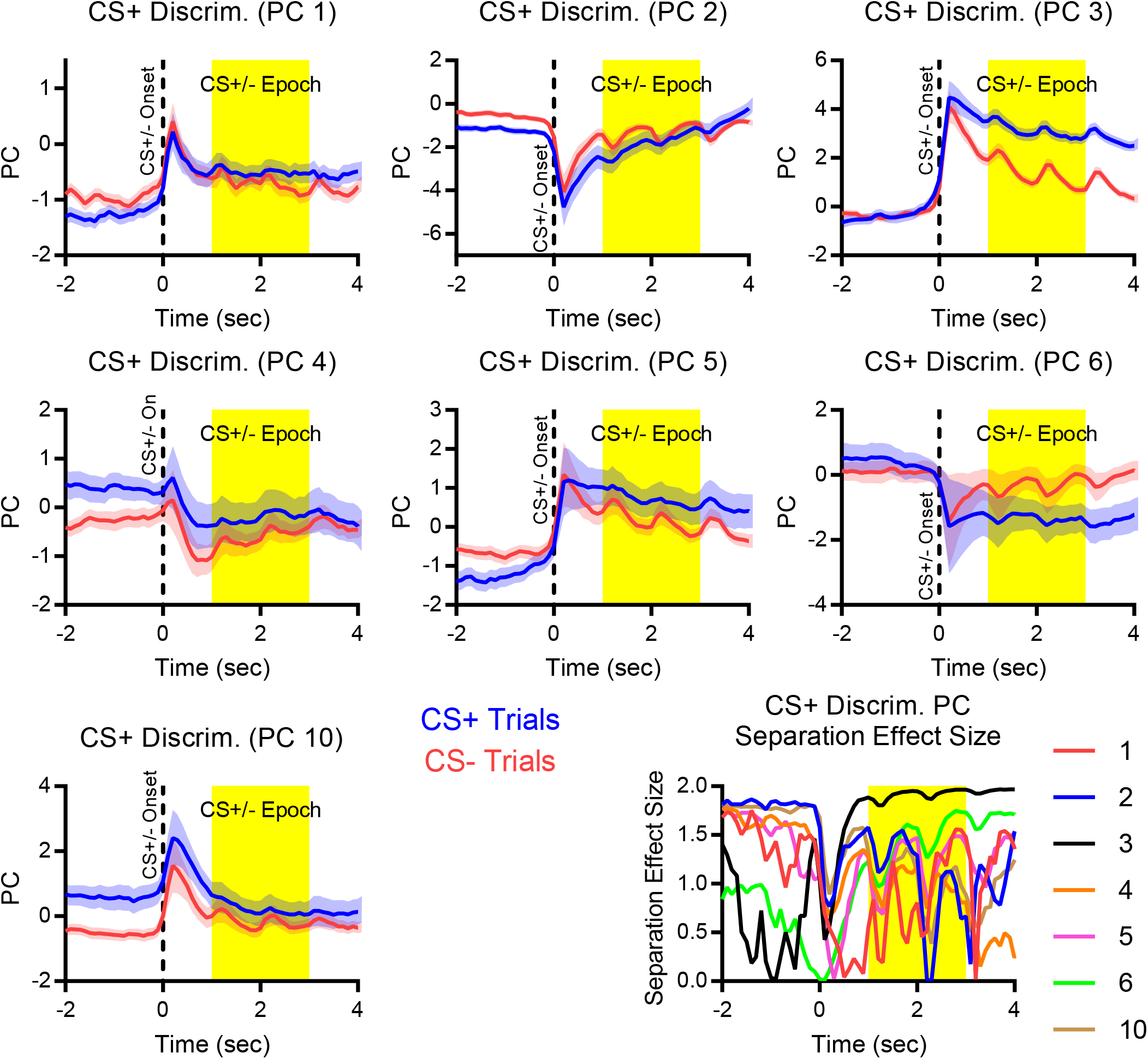
All stable PCs (all neurons) averaged over CS+ and CS-trials (mean +/-std). CS+ vs. CS-representation strength was assessed by comparing the CS+ and CS-PC values during the CS+/-epoch (yellow region). PC separation was weakest for CS discrimination, with PCs 3 and 6 showing the largest separation.

**Supplemental Figure 7:**
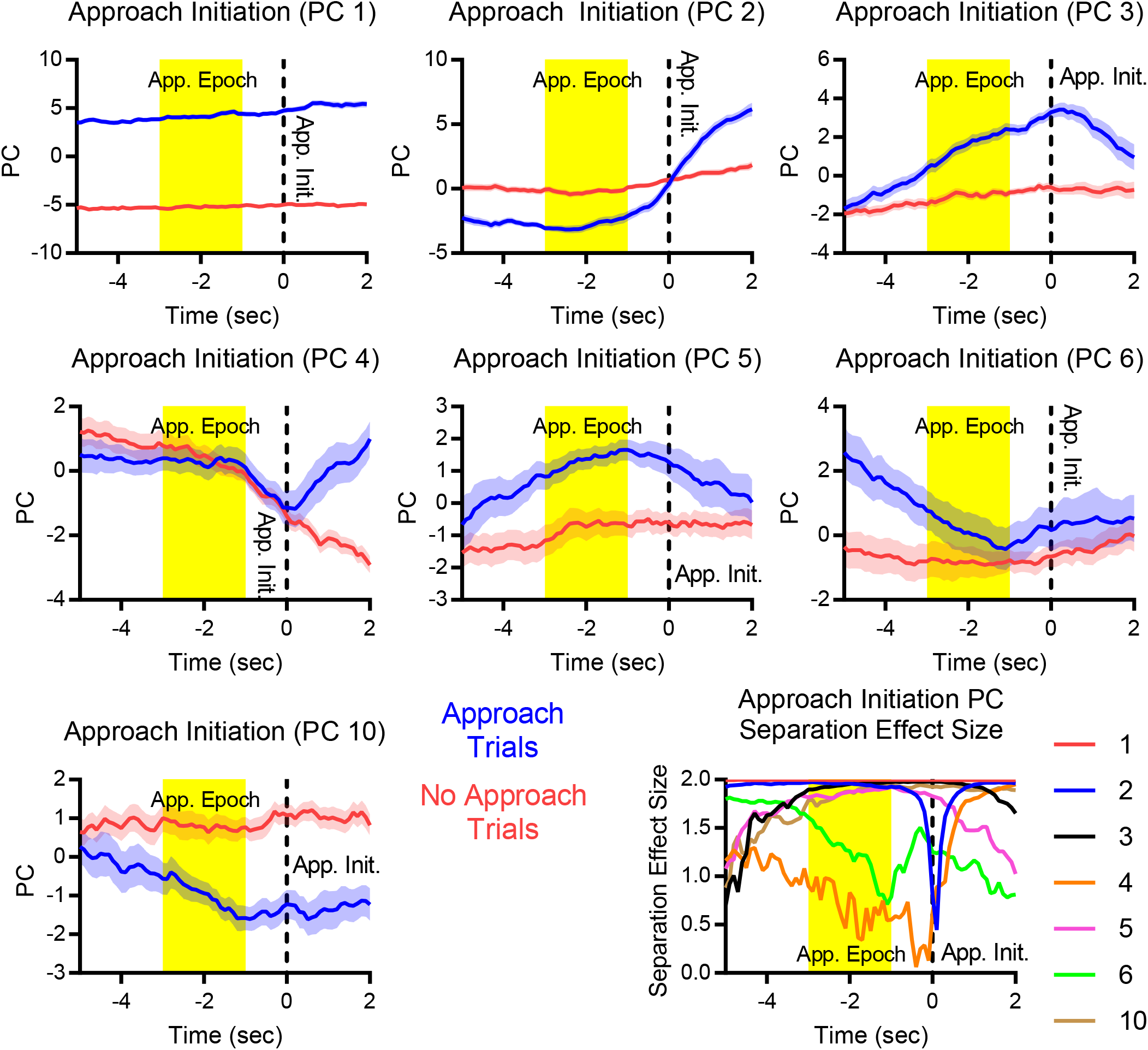
All stable PCs (all neurons) averaged over approach and no approach trials time locked to the approach initiation time point found using the change point in the within trial likelihood to approach (mean +/-std). Approach initiation representation strength was assessed by comparing PC trajectories on approach and no approach trials during the approach initiation time window (yellow epoch). Most PCs showed robust separation during the approach initiation epoch, many with distinct dynamics over the time window shown.

**Supplemental Figure 8:**
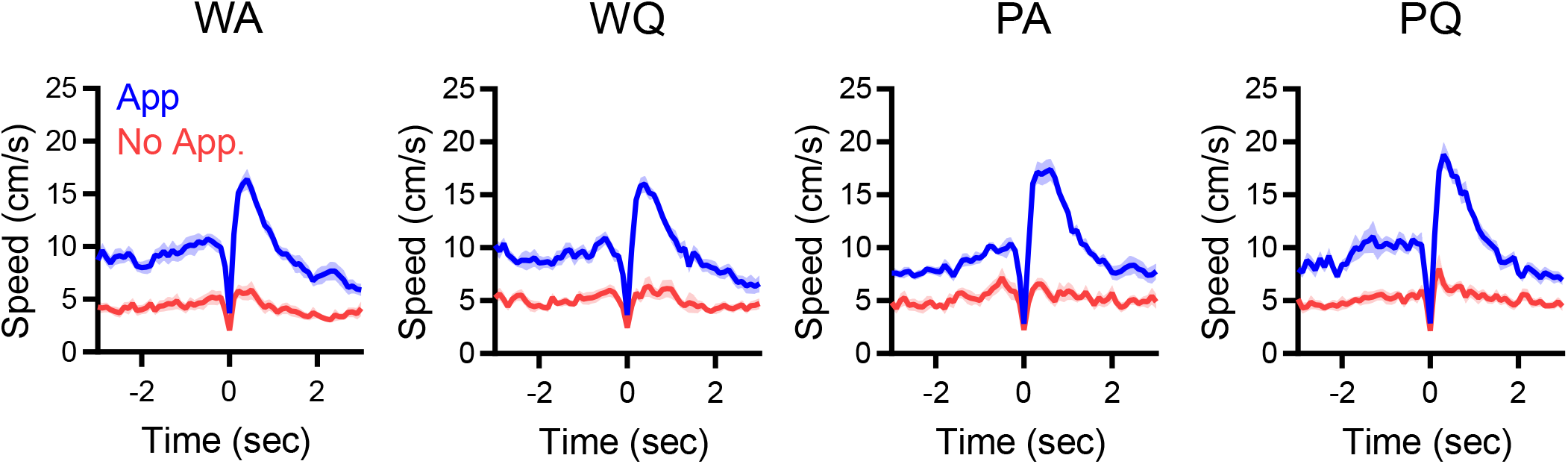
Speed profiles time locked to approach initiation for all sessions. Approach trials tended to have higher speed before approach initiation and a large increase in speed following approach initiation. W: Wistar. P: P rat. A: alcohol. Q: quinine.

**Supplemental Figure 9:**
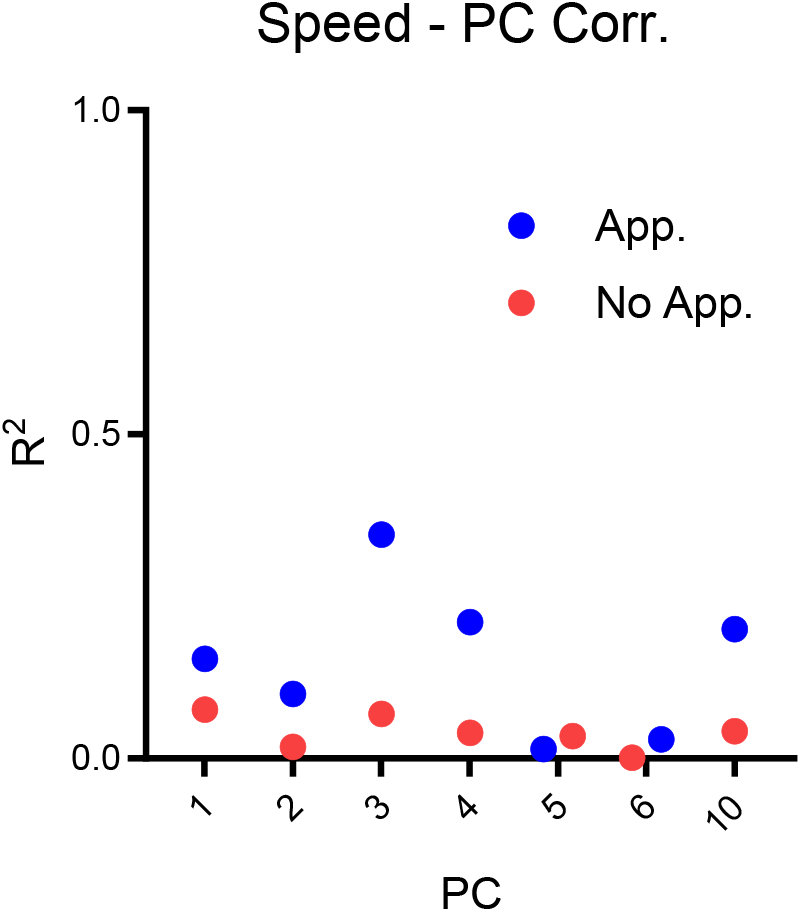
PC scores were weakly correlated with animal speed. From 3 seconds before to 2 seconds after approach initiation, the correlation between the speed (Supplemental Figure 8) and the PC scores (Supplemental Figure 7) was low. Furthermore, no PCs exhibited sharp changes near the approach initiation, despite the presence of characteristic decreases in speed near approach initiation. This indicates that these PCs were not simply movement artifacts or represented only movement.

**Supplemental Figure 10:**
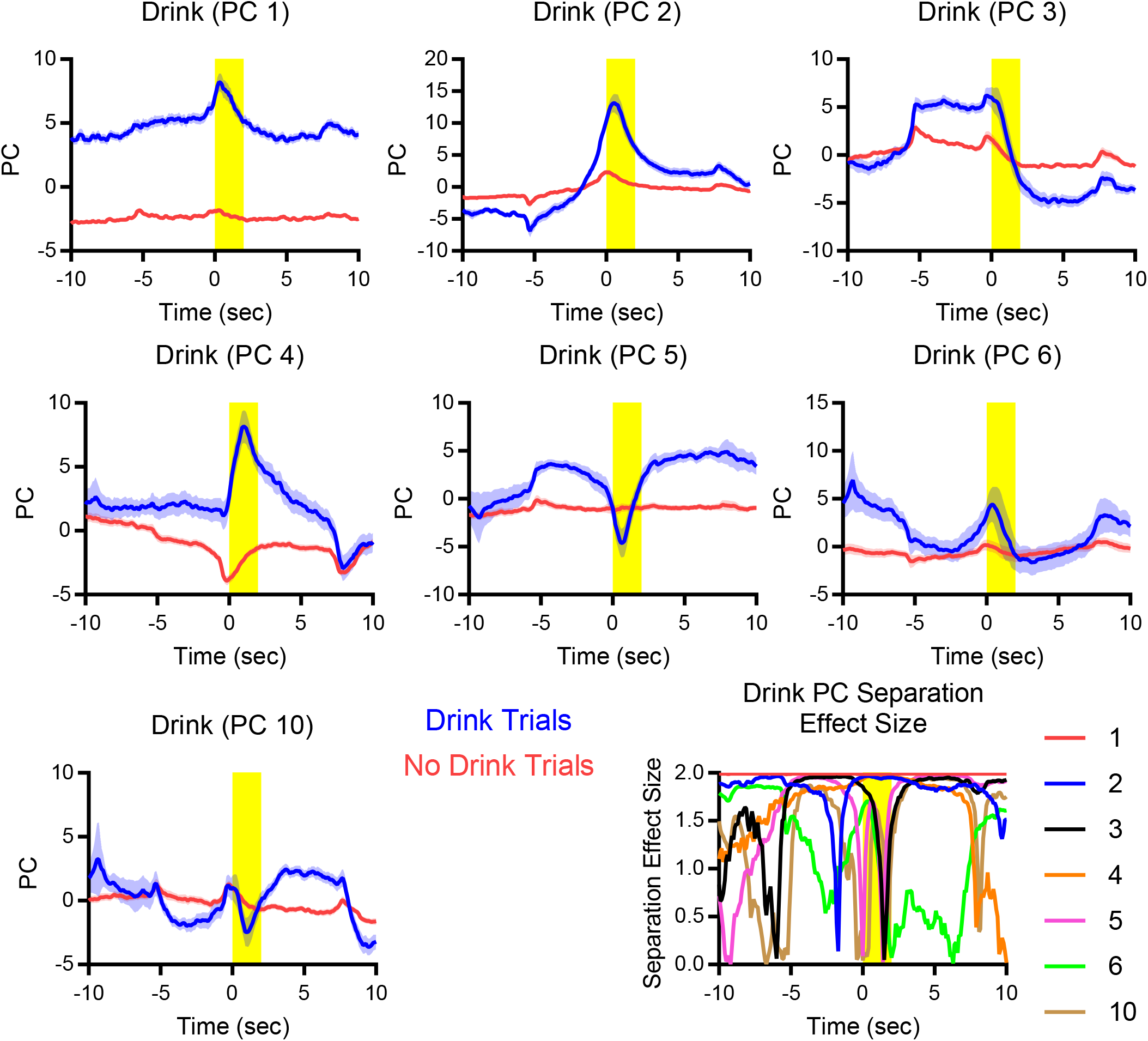
All stable PCs (all neurons used) averaged over drink and no drink trials time locked to the time the animal arrived at the sipper (mean +/-std). Drink encoding was assessed by comparing PC trajectories on drink and no drink trials during the drink time window (yellow epoch). Many PCs showed robust separation during the drink epoch, many with distinct dynamics over the time window shown.

**Supplemental Figure 11:**
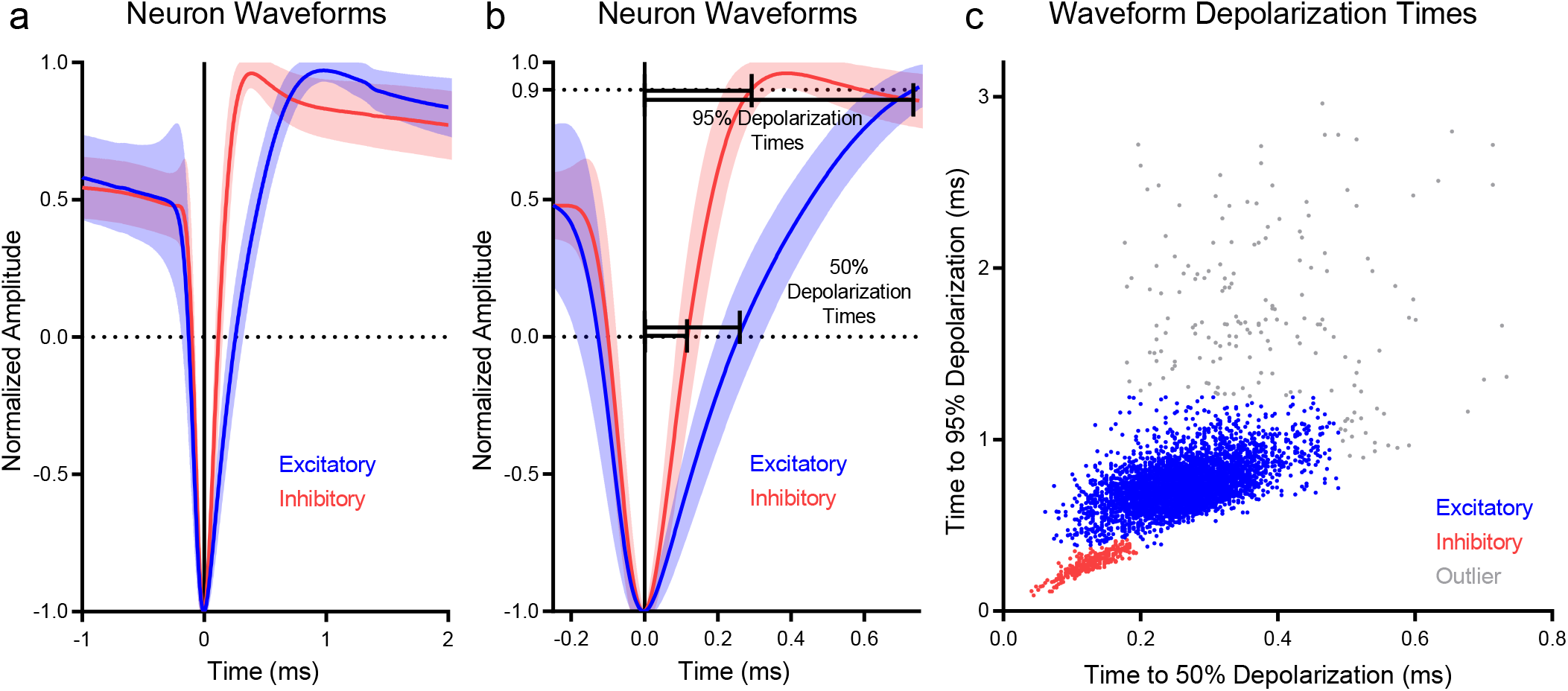
Excitatory and inhibitory neuron classification by waveform. **a**, Average neuron waveforms (taken from electrode with largest amplitude) were scaled to range from -1 to 1. **b**, To measure how fast the action potential depolarized, we assessed the time it took the average waveform to reach 50% (0) and 95% (0.9) of the maximum depolarization. **c**, When these times were plotted, two clusters and outliers were apparent. Manual cluster was performed to label neurons as excitatory or inhibitory.

**Supplemental Figure 12:**
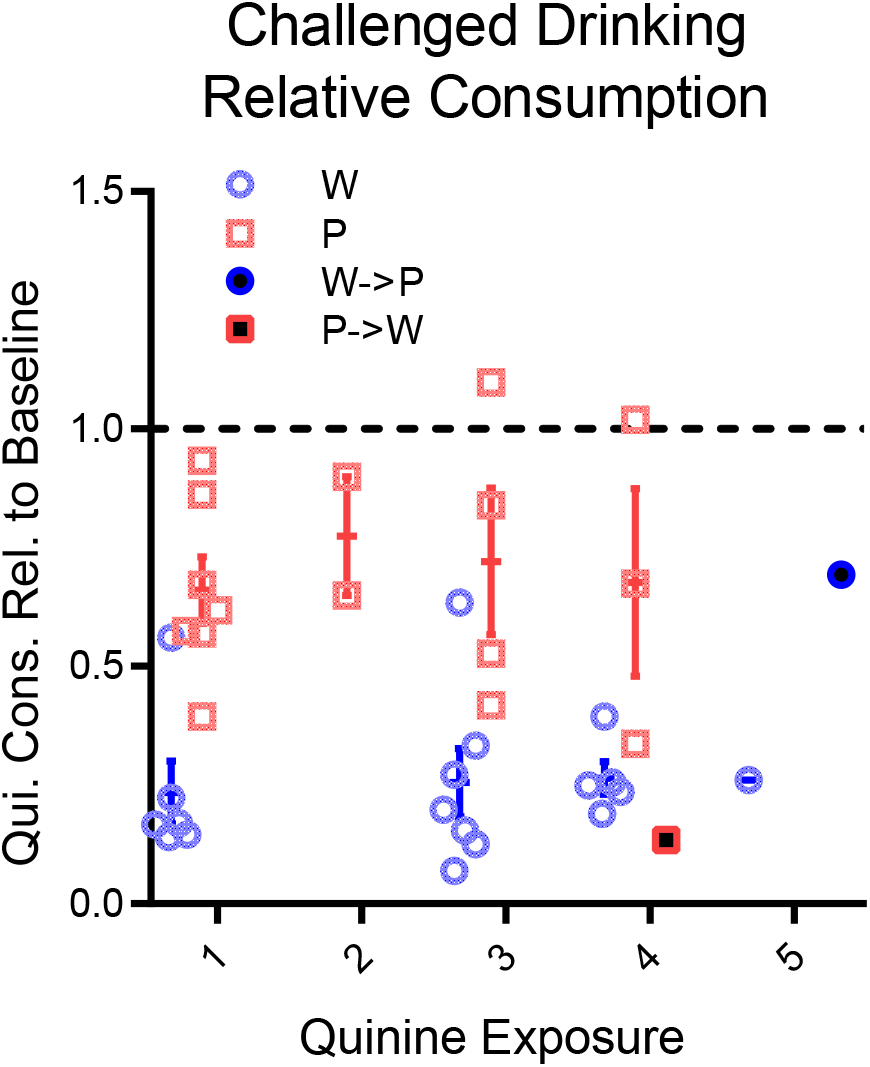
Challenged drinking behavior was largely consistent across later sessions. Relative consumption between challenged drinking (quinine) and baseline consumption on the previous day remained relatively unchanged across additional quinine exposures. Sample sizes varied across quinine exposures due to ordering of experimental sessions. For all Wistars, the second quinine exposure was the free access quinine test. One Wistar and one P rat were switched between groups for analysis of electrophysiology (see Supplemental Fig. 19).

**Supplemental Figure 13:**
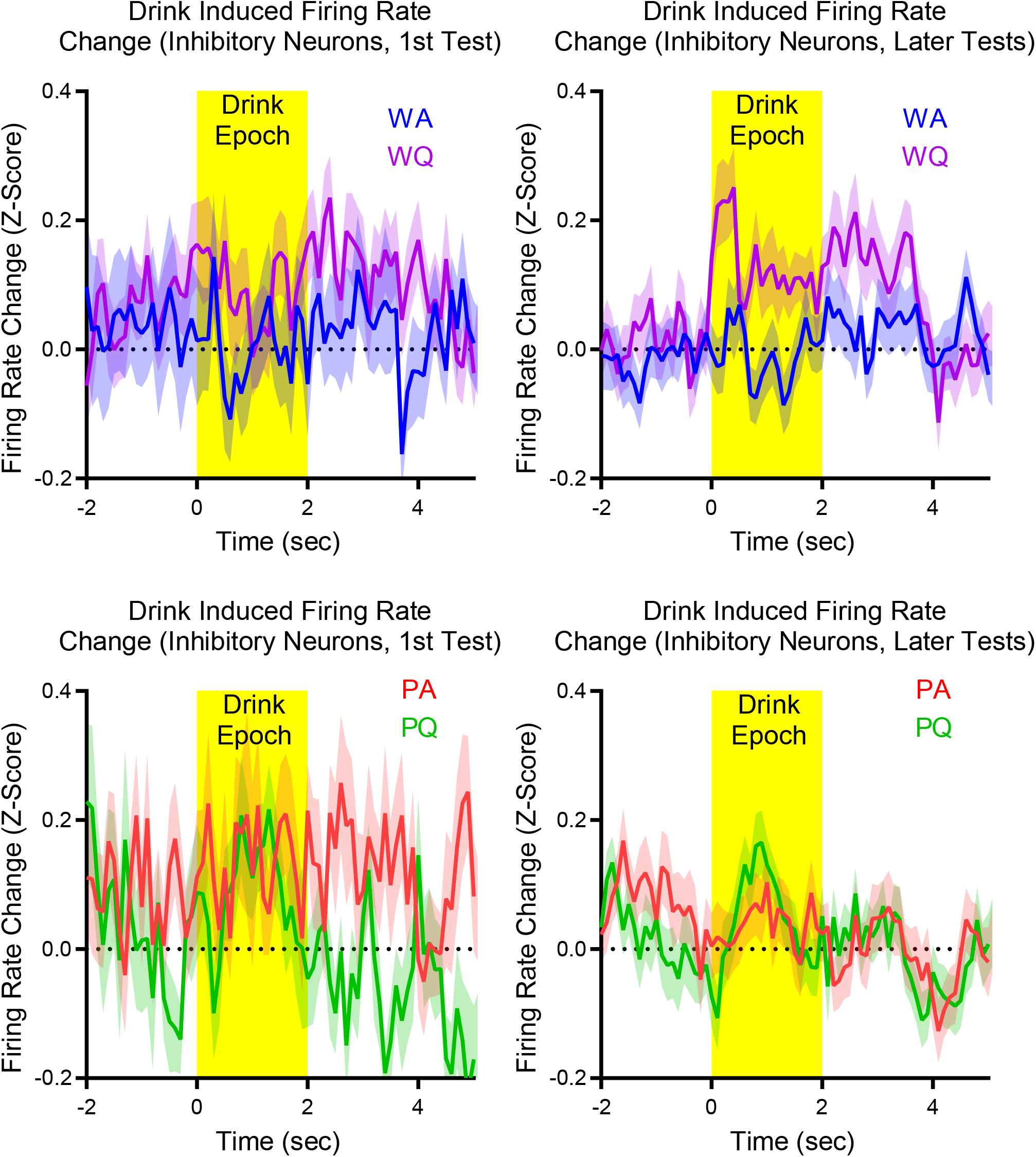
Inhibitory neuron firing rate responses to drink trials. Significant differences were found for the mean firing rate change during the drink epoch in Wistars during later compulsive drinking tests.

**Supplemental Figure 14:**
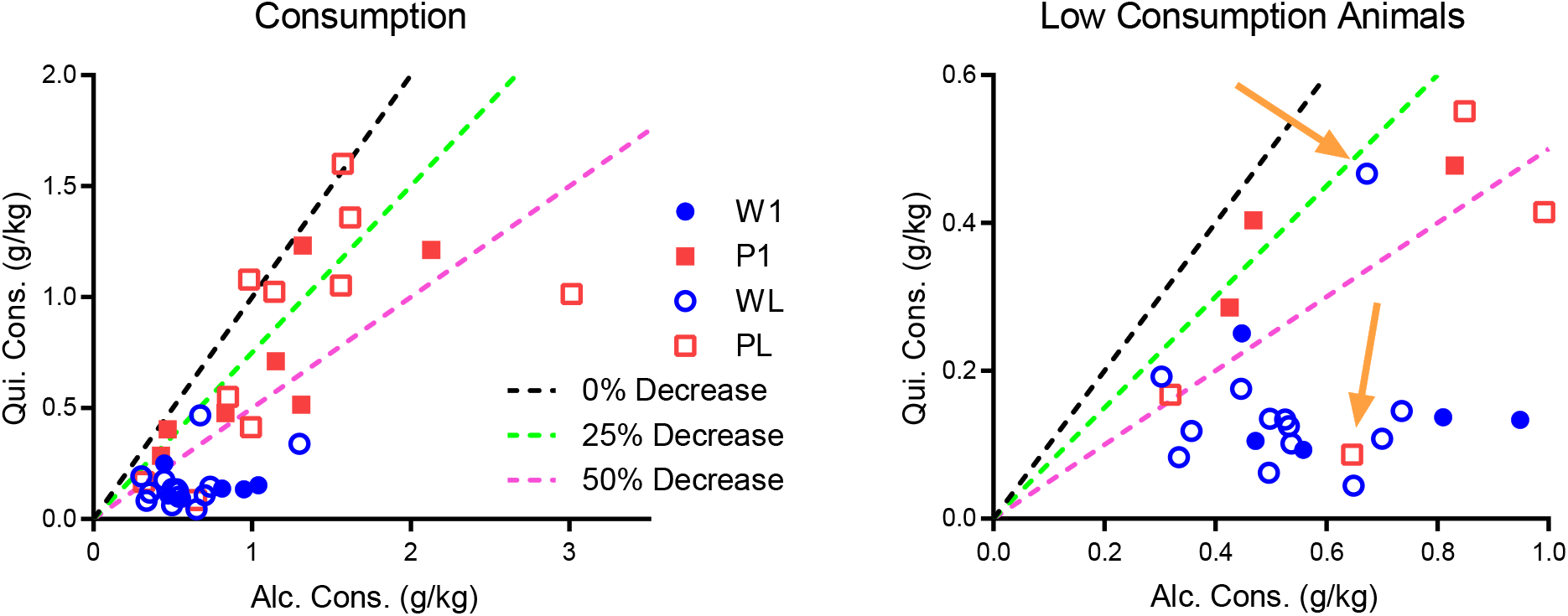
For later recordings, in the PCA, two subjects were switched between groups based on alcohol and quinine consumption (orange arrows). One Wistar exhibited a small drop in consumption when quinine was added. Also, one P rat exhibited a large drop in consumption when quinine was added. Animals with lower alcohol consumption values (below 0.5 g/kg), were not switched due to the relative magnitude of the consumption measurement error.

**Supplemental Figure 15:**
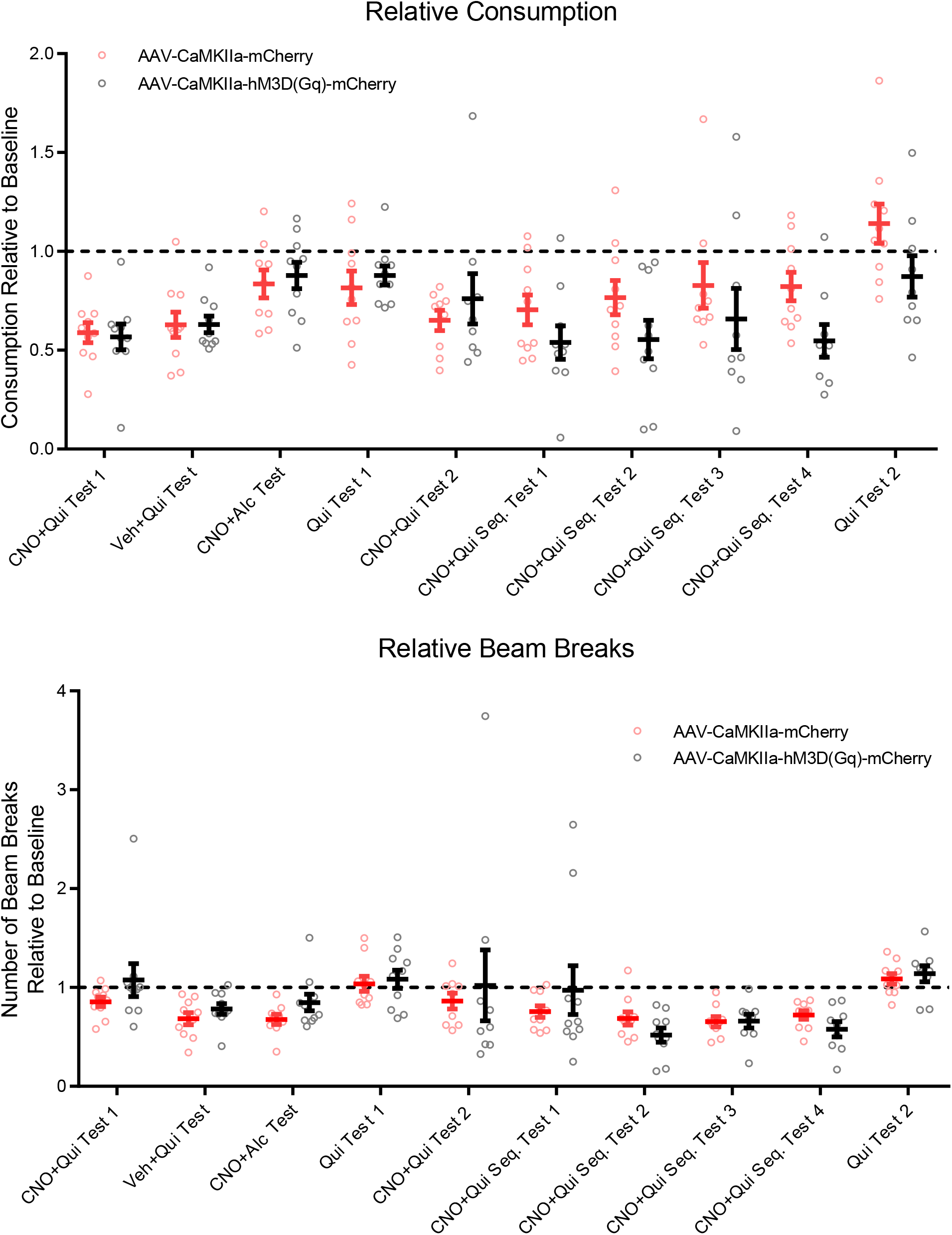
Behavioral tests in DREADD and control P rats. Tests were performed sequentially from left to right with regular 2CAP session interspersed. All animals underwent a CNO+quinine test first. Half of the animals then underwent a vehicle+quinine test, a regular quinine test with no injection, then a CNO+alcohol test. The other animals underwent the same tests, but in reversed order. All animals then underwent a CNO+quinine test, followed by a sequence of four CNO+quinine tests for four days (one each day, no regular 2CAP sessions between). Finally, all animals underwent a quinine test with no injection. No differences were observed between number of beam breaks between control and DREADD animals the two groups of tests discussed in the main text or as post-doc comparisons for individual tests. (Mean +/-sem)

**Supplemental Figure 16:**
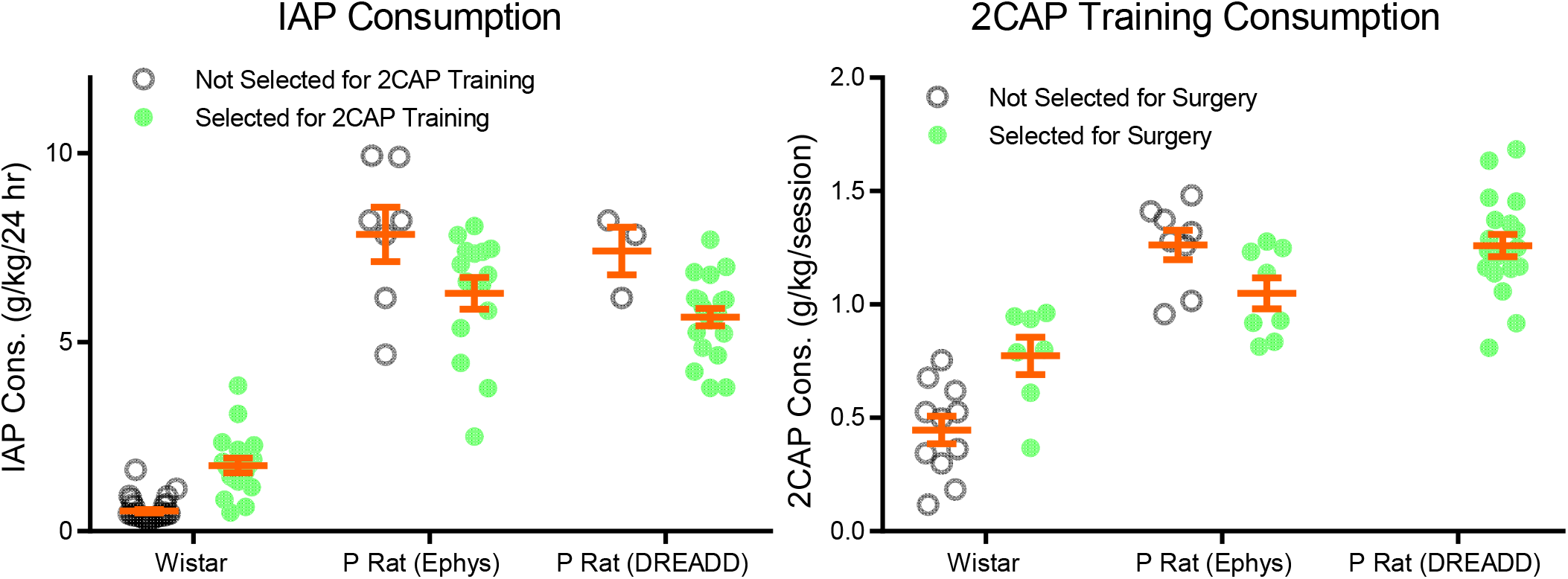
Consumption results for animals during IAP and 2CAP training. Animals were primarily selected based on average consumption levels. However, this was not always possible due to the sizes of the cohorts. Also, some animals were not selected due to health concerns. In general, higher drinking Wistars and lower drinking P rats were selected.

**Supplemental Figure 17:**
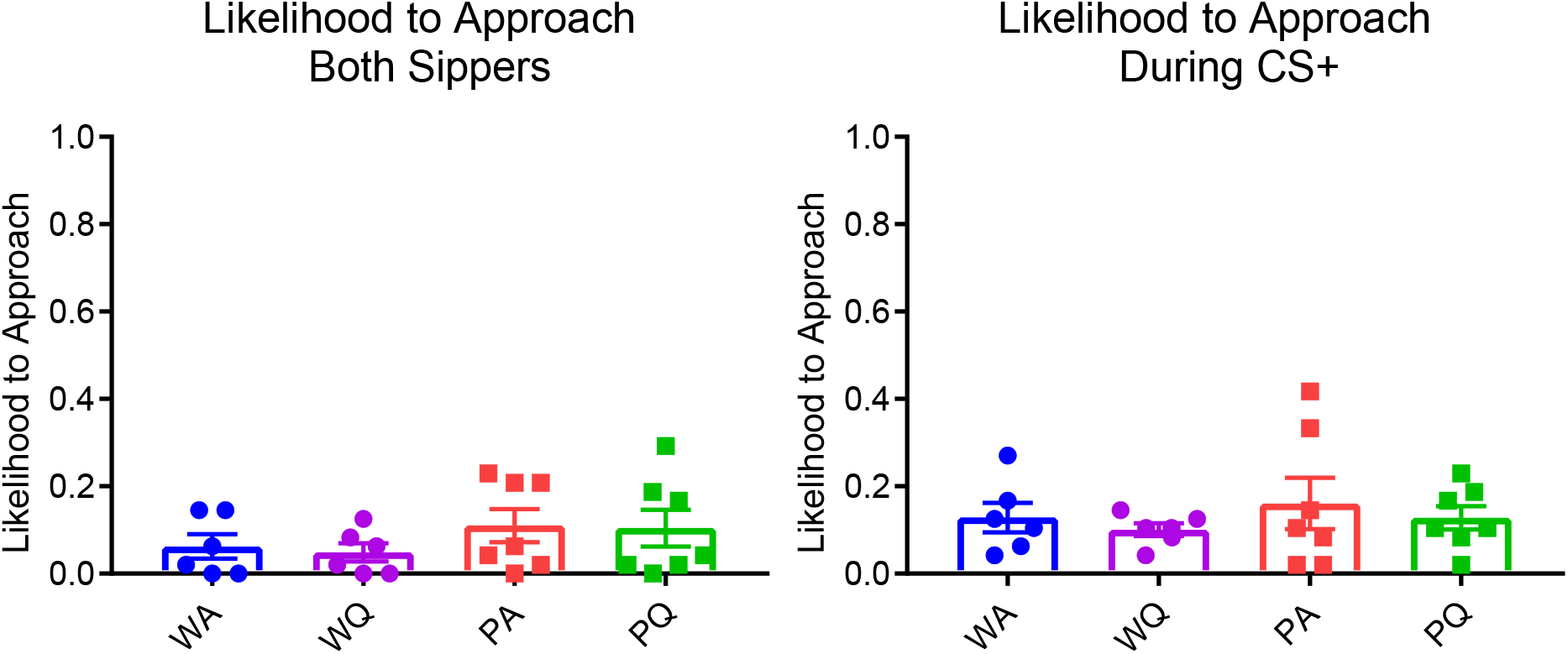
Animals were unlikely to approach both sippers during access (i.e., correct from the incorrect sipper to the correct sipper) or approach the correct sipper during the CS+. No significant differences were observed between groups.

**Supplemental Figure 18:**
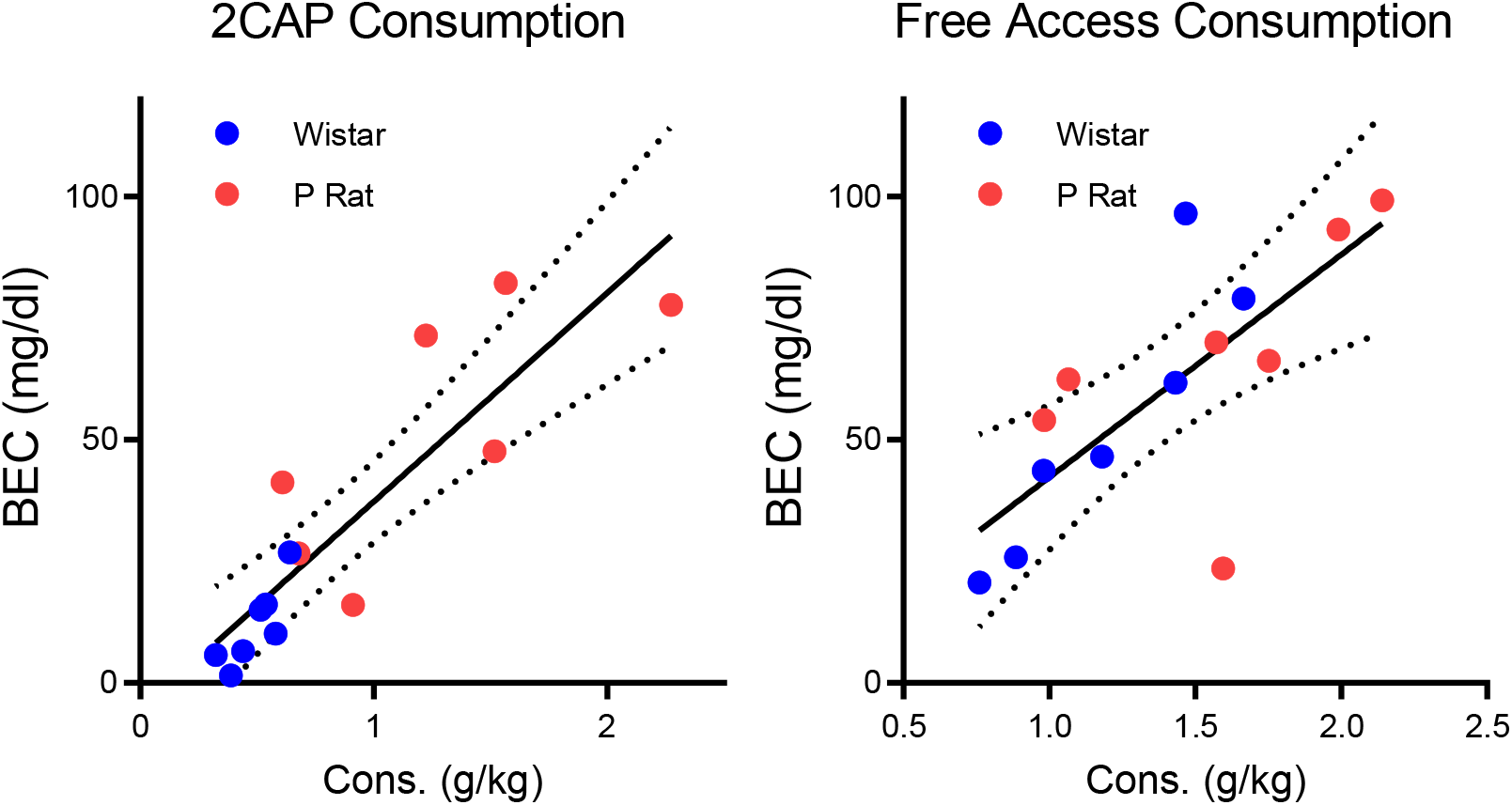
Blood ethanol concentration measurements for regular 2CAP sessions and free access sessions. During the free access session, the animal was given unlimited access to both sippers for the same time period as a regular 2CAP session. The free access sessions and the regular 2CAP sessions shown here were not part of the rest of the analysis. A fit of the regular 2CAP intake vs. BEC for all animals found a significant slope (mean +/-SE: 42.94 +/-6.87, F(1,12) = 39.02, p < 0.0001, R^2^ = 0.76). Similarly, a fit of the free access intake vs. BEC for all animals also found a significant slope (mean +/-SE: 45.78 +/-12.09, F(1,12) = 14.34, p = 0.0026, R^2^ = 0.54).

**Supplemental Figure 19:**
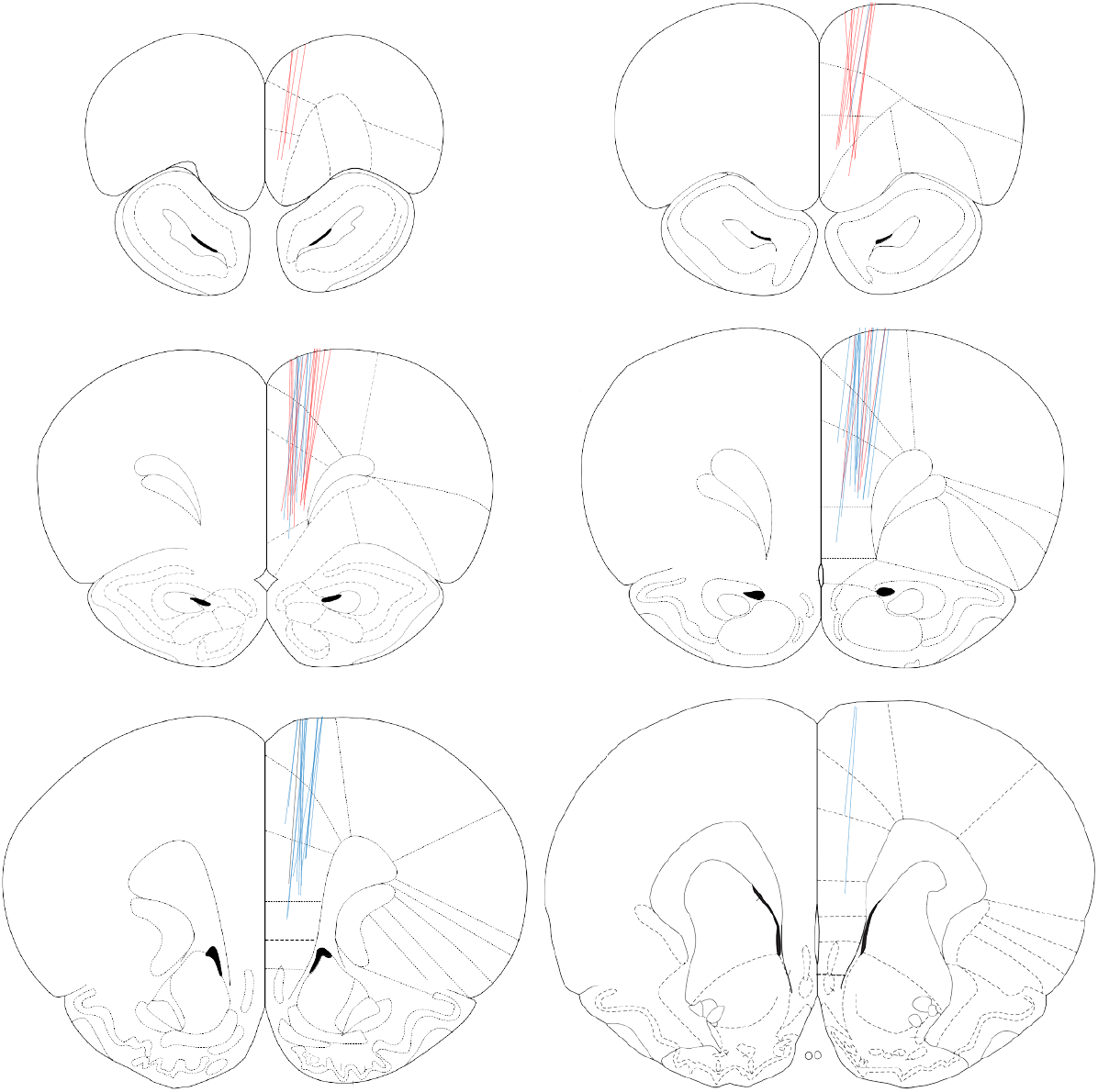
Placement tracks for all Wistars (blue) and P rats (red).

**Supplemental Figure 20:**
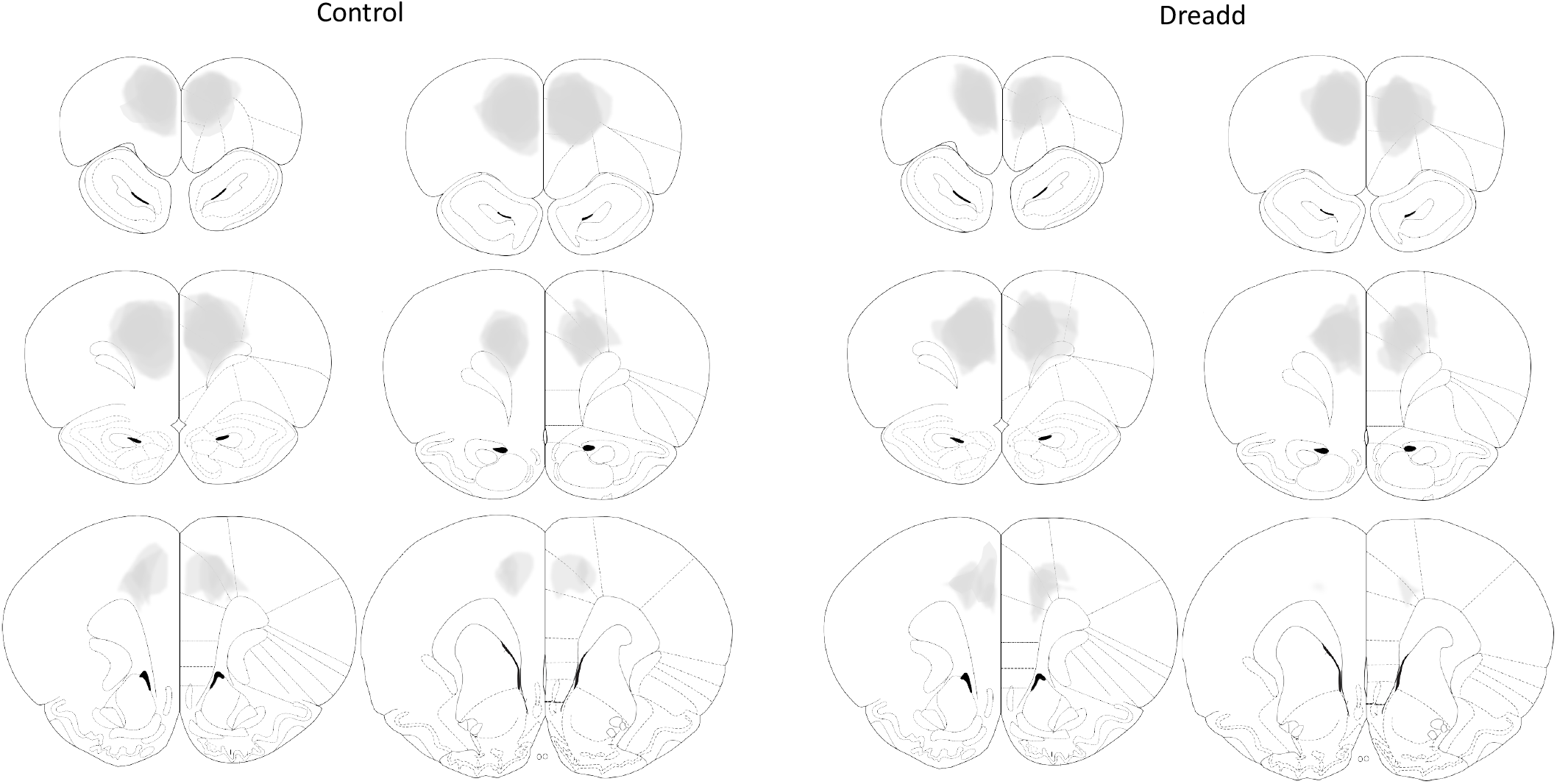
Virus expression for control and DREADD animals. Expression was verified for all animals.

**Supplemental Figure 21:**
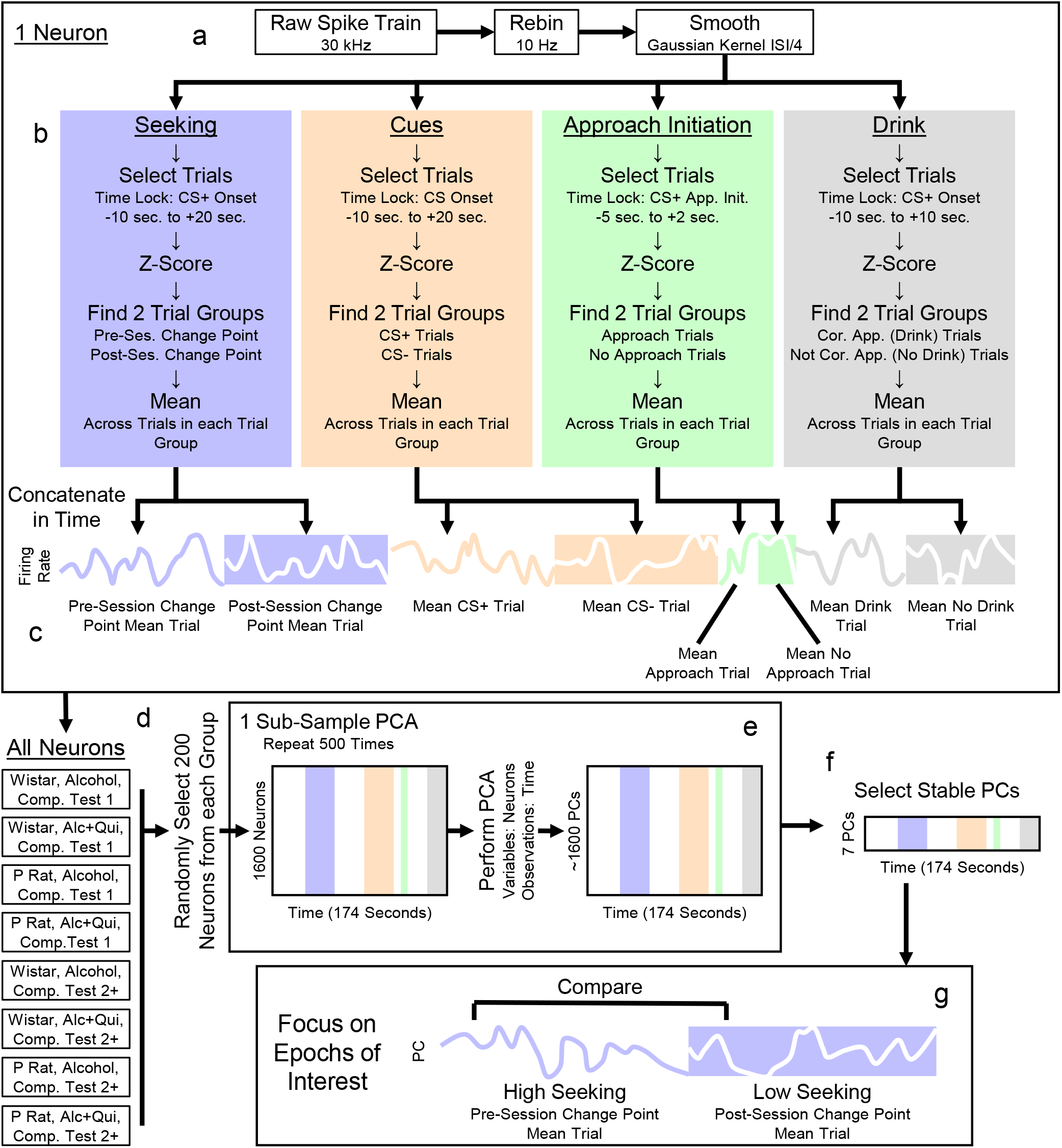
PCA workflow. a) Raw data was recorded and spike sorted at 30 kHz, then downsampled to 10 Hz, then smoothed with a Gaussian kernel (standard deviation 25% of mean ISI separation for each neuron). b) For each segment of the decision-making process of interest, trials were selected based on a time lock point and a time window. The data were z-scored across all trials of interest. The trials were divided into two groups and means were taken across trials in each group. c) All mean trials of interest were concatenated for each neuron. d) Neurons were group by experimental conditions (8 groups: P rat vs. Wistar, alcohol vs. alcohol+quinine, Compulsion Test 1 vs. Compulsion Test 2+). e) Principal Component Analysis (PCA) was conducted using multiple subsamples to control for neuron yield in each experimental group. In each PCA iteration, 200 neurons from each experimental group were randomly selected. PCA was performed using neurons as variables and time as observations to assess neural population firing patterns through trials of interest. f) Stable PCs were identified by comparing mean signal variance to variance across PCA iterations. g) Specific comparisons were made between PCs to assess encoding of various decision-making related variables, such as high seeking trials vs. low seeking trials.

